# Extended toolboxes enable efficient biosynthesis of valuable chemicals directly from CO_2_ in fast-growing *Synechococcus* sp. PCC 11901

**DOI:** 10.1101/2023.08.23.554402

**Authors:** Tong Zhang, Shubin Li, Lei Chen, Tao Sun, Weiwen Zhang

## Abstract

CO_2_ recycle is crucial to the global carbon neutrality. Though cyanobacteria are known to be photoautotrophic cell factories capable of converting CO_2_ into valuable chemicals, their slower growth rate and lower biomass accumulation compared to those of the heterotrophic organisms significantly restrict their application at commercial scale. The newly discovered marine cyanobacterium, Synechococcus sp. PCC 11901 (hereafter PCC 11901) offers several advantages like rapid growth, high biomass and high salinity tolerance, and could become a new generation of cyanobacterial chassis. To promote its application, in this study we developed genetic toolboxes applicable to PCC11901. First, a cobalamin (V_B12_)-independent chassis was constructed, allowing for its cheaper cultivation. Second, genome copy numbers and transformation methods were respectively measured and optimized. The 14 neutral sites were identified and characterized within the genome PCC 11901, providing locations for genetic integration of exogenous cassettes. Subsequently, libraries were developed, reaching an expression range of approximately 800 folds for constitutive promoters and an induction fold of up to approximately 400 for inducible promotor, respectively. As a proof of concept of its utilization, we engineered the synthetic pathways of glucosylglycerol (GG) into PCC 11901 using the established toolboxes, yielding 590.41 ± 21.48 mg/L for GG production. Notably, we found the cobalamin-independent PCC 11901 chassis exhibited superior self-sedimentation ability compared to the wild-type chassis. Our work here made it possible to develop the fast-growing PCC 11901 as efficient carbon-neutral cell factory in the future.

## 1. Introduction

The global industrialization process has greatly exacerbated the emission of CO_2_, resulting in global warming and extreme weather conditions. Over the past decades, the burning of fossil fuels has led to a steady increase in CO_2_ emissions, rising from an average of 11 billion tons per year in the 1960s to 35 billion tons per year in the 2010s [1]. Consequently, there has been a growing interest in recycling CO_2_ to back into high-value chemicals [2]. Cyanobacteria, which are the only prokaryotes capable of oxygenic photosynthesis, have garnered attention due to their ability to harness solar energy to fix CO_2_ and convert it into biomass and metabolites. Notably, the average photosynthetic conversion rate and growth rate are comparable to microalgae and surpass those of terrestrial plants [3, 4]. In recent years, genetic modifications have been successfully implemented in cyanobacteria to synthesize a wide range of non-natural compounds, including commodity chemicals, biofuels, and high-value products [5–7]. These studies demonstrate the numerous advantages of cyanobacteria as “light-driven autotrophic cell factories” for CO_2_ fixation and utilization.

For many years, several dozens of valuable chemicals like 3-hydroxypropionic acid [8], *myo*-inositol [9], astaxanthin [10], α-farnesene [11], limonene [12], sucrose [13], and ethylene [14] have been synthesized in various cyanobacterial chassis. Despite these successful efforts, the industrial application of cyanobacteria still faces obstacles such as low productivity and poor robustness. The problem is mainly due to their slow growth rate and low biomass accumulation of cyanobacteria. For example, cyanobacteria are typically with a doubling time of approximate 6.6 h for *Synechocystis* sp. PCC 6803 (PCC 6803) and 4.9 h for *Synechococcus elongatus* PCC 7942 (PCC 7942), respectively, along with a lower biomass accumulation (average ∼5 g·L^-1^ in 10 days) [15]. To address these issues, one of the key strategies involves discovery and utilization of more promising and efficient cyanobacterial chassis. For example, Wendt et al. [16] identified and characterized *Synechococcus elongatus* UTEX 2973 (UTEX 2973), which is with a doubling time of approximately 1.5 hours. This species has emerged as a strong candidate for large-scale applications in recent study [17]. Similarly, Włodarczyk et al. [18] discovered that *Synechococcus* sp. PCC 11901 (hereafter PCC 11901) exhibited higher biomass production even under high-light intensities (up to 750 µmol photons m^−2^ s^−1^), with a multiplication time as short as 2 hours. Moreover, PCC 11901 is highly tolerant to extreme temperatures (up to 43°C), high-light irradiance, and salinity, resulting in greater biomass accumulation (up to ∼33 gDW/L) under its optimal growth condition. Notably, the synthesis of free fatty acids (FFA) in PCC 11901 chassis yielded 1.54 g/L, which is comparable to that achieved in engineered *Escherichia* coli with similar genetic manipulation. Therefore, PCC 11901 represents one of the most promising candidates as an industrial biotechnology platform for the sustainable production of chemical products. However, the available genetic toolboxes in PCC 11901 are significantly limited currently compared to other model cyanobacterial species. This limitation severely restricts further research and the application of synthetic biology approaches in this species.

To address the aforementioned issues, we aimed to extend the toolboxes available for PCC 11901 in this study (**Fig. 1**). First, we developed mutant strains that exhibits normal growth independent on expensive cobalamin. Then the genome copy numbers and transformation methods were explored. In addition, we identified and characterized some new neutral sites. Moreover, we established libraries of constitutive and inducible promoters separately, allowing for a wide range of gene expression regulation. Utilizing these tools, we constructed synthetic pathways for glucosylglycerol (GG), marking the first successful synthesis of this compound in PCC 11901. Finally, we investigated the self-sedimentation capability of the engineered bacteria. The synthetic biology tools and chemical synthesis pathways proposed in this study are crucial for establishing PCC 11901 as a “photosynthetic cell factory” of industrial scale in the ne future.

**Fig. 1.**
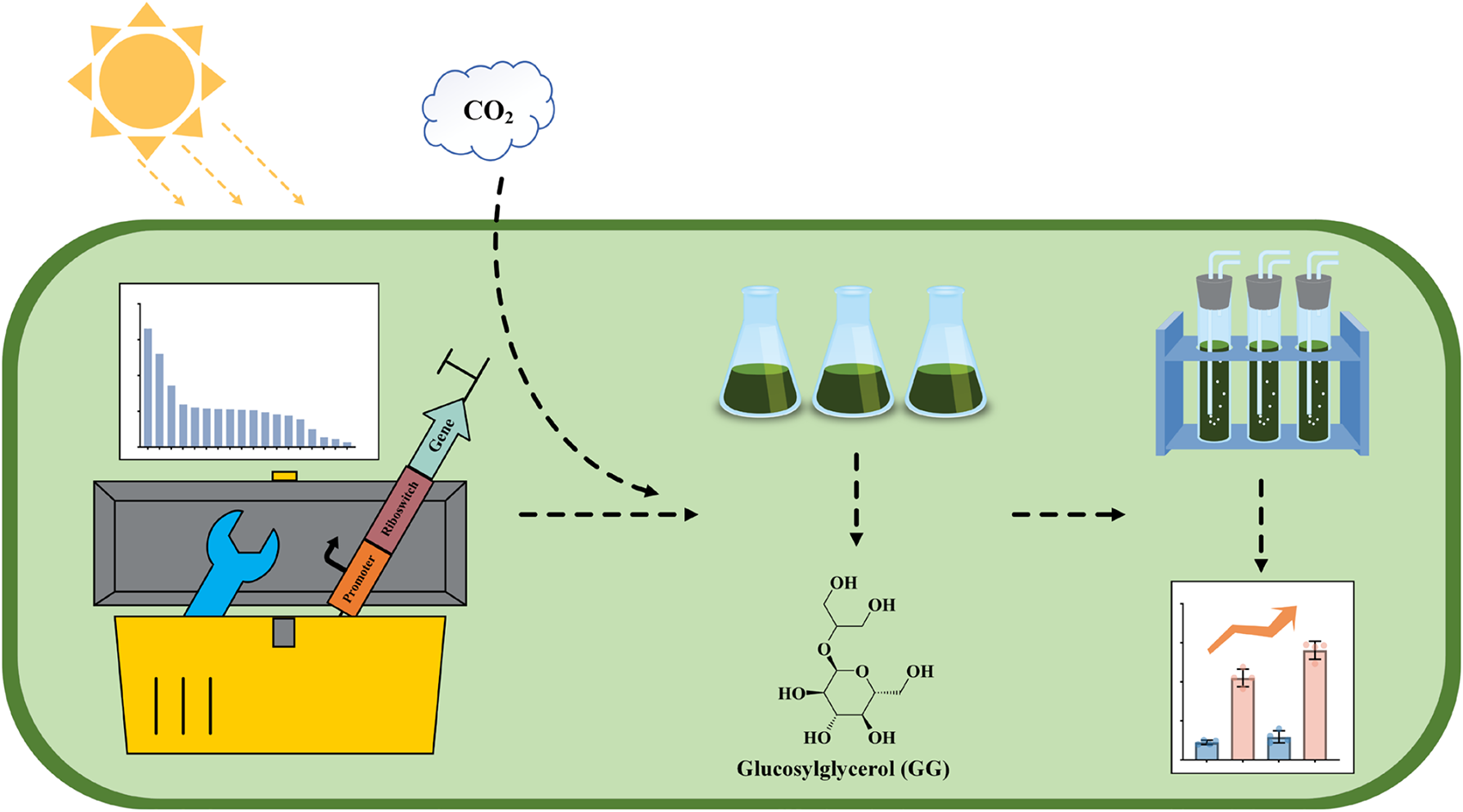
Schematic of this study.

## 2. Material and methods

### 2.1 Bacterial growth conditions and strains construction

The wild-type strain of PCC 11901 (WT) was kindly provided by Prof. Fei Tao from Shanghai Jiao Tong University. Both the WT and genetically engineered strains were cultivated in MAD medium (pH 8.1) at a constant light intensity of approximately 100 μmol photons m^-2^ s^-1^. The cultivation was carried out at 37°C with a shaking incubator (HNYC-202T, Honour, Tianjin, China) or on MAD agar medium in an illuminated incubator (SPX-250B-G, Boxun, Shanghai, China). To ensure the stability of the genetically engineered strains, suitable antibiotics were added as needed (spectinomycin, 20 μg/mL; kanamycin, 50 μg/mL) [18, 19]. A high-density culture of PCC 11901 was conducted in an incubator (SPX-350, Zhongyiguoke, Beijing, China) at a light intensity of 600 μmol photons m^-2^ s^-1^ and a temperature of 37°C bubbled with 10% (v/v) CO_2_. For the cultivation of *E. coli* strains, LB agar medium or liquid medium supplemented with appropriate antibiotics was used. The incubation was carried out at 37°C either in an incubator (HNY-100B, Honour, Tianjin, China) or a shaking incubator (HNY-100B, China) at 200 rpm. To maintain the stability of the plasmids, antibiotics were added at the following working concentrations: 100 μg/mL for spectinomycin, 100 μg/mL for ampicillin, and 50 μg/mL for kanamycin. The absorbance of PCC 11901 and *E. coli* suspensions was respectively measured at OD_750 nm_ and OD_600 nm_, using an ELx808 absorbance microporous plate (BioTek, Winooski, VT, USA).

The construction and amplification of plasmids were conducted using the DH5α strain of *E. coli*. The primers utilized in this study were synthesized by GENEWIZ Inc. (Suzhou, China). **Table S1** displays the genetic elements employed. All strains employed or generated in this study can be found in **Table 1**. Verification of all transformants was carried out through colony PCR and DNA sequencing.

**Table 1.**
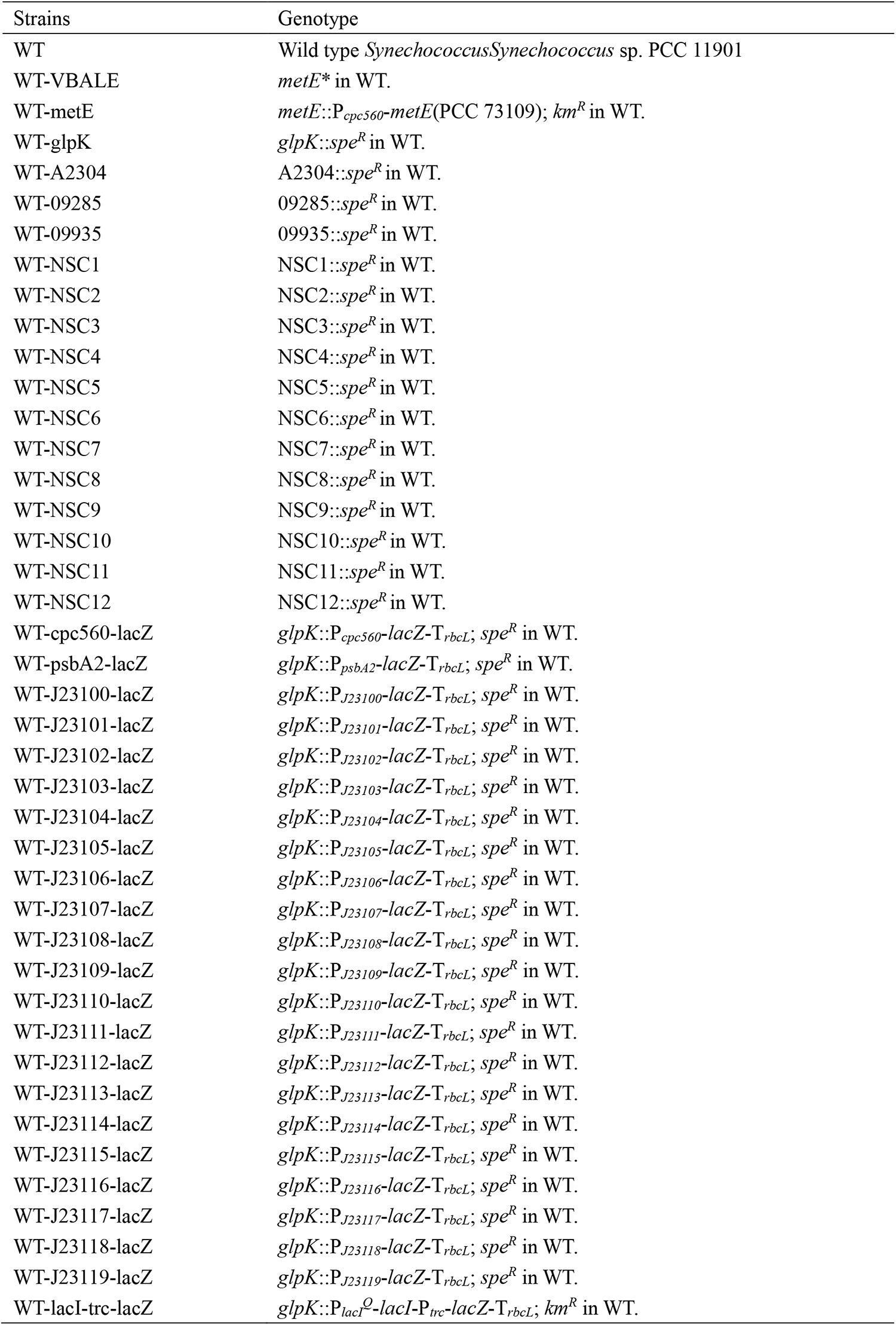
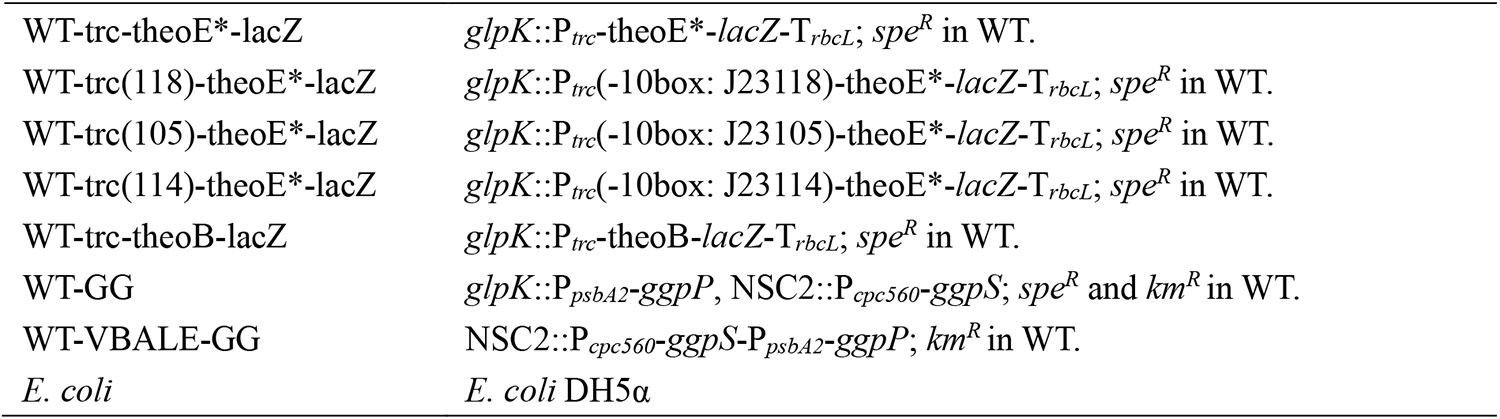
Strains constructed in this study.

| Strains | Genotype |
| --- | --- |
| WT | Wild type *Synechococcus* *Synechococcus* sp. PCC 11901 |
| WT-VBALE | *metE* * in WT. |
| WT-metE | *metE*::*P<sub>cpc560</sub>-metE*(PCC 73109); *km<sup>R</sup>* in WT. |
| WT-glpK | *glpK*::*spe<sup>R</sup>* in WT. |
| WT-A2304 | A2304::*spe<sup>R</sup>* in WT. |
| WT-09285 | 09285::*spe<sup>R</sup>* in WT. |
| WT-09935 | 09935::*spe<sup>R</sup>* in WT. |
| WT-NSC1 | NSC1::*spe<sup>R</sup>* in WT. |
| WT-NSC2 | NSC2::*spe<sup>R</sup>* in WT. |
| WT-NSC3 | NSC3::*spe<sup>R</sup>* in WT. |
| WT-NSC4 | NSC4::*spe<sup>R</sup>* in WT. |
| WT-NSC5 | NSC5::*spe<sup>R</sup>* in WT. |
| WT-NSC6 | NSC6::*spe<sup>R</sup>* in WT. |
| WT-NSC7 | NSC7::*spe<sup>R</sup>* in WT. |
| WT-NSC8 | NSC8::*spe<sup>R</sup>* in WT. |
| WT-NSC9 | NSC9::*spe<sup>R</sup>* in WT. |
| WT-NSC10 | NSC10::*spe<sup>R</sup>* in WT. |
| WT-NSC11 | NSC11::*spe<sup>R</sup>* in WT. |
| WT-NSC12 | NSC12::*spe<sup>R</sup>* in WT. |
| WT-cpc560-lacZ | *glpK*::*P<sub>cpc560</sub>-lacZ-T<sub>rbcL</sub>*; *spe<sup>R</sup>* in WT. |
| WT-psbA2-lacZ | *glpK*::*P<sub>psbA2</sub>-lacZ-T<sub>rbcL</sub>*; *spe<sup>R</sup>* in WT. |
| WT-J23100-lacZ | *glpK*::*P<sub>J23100</sub>-lacZ-T<sub>rbcL</sub>*; *spe<sup>R</sup>* in WT. |
| WT-J23101-lacZ | *glpK*::*P<sub>J23101</sub>-lacZ-T<sub>rbcL</sub>*; *spe<sup>R</sup>* in WT. |
| WT-J23102-lacZ | *glpK*::*P<sub>J23102</sub>-lacZ-T<sub>rbcL</sub>*; *spe<sup>R</sup>* in WT. |
| WT-J23103-lacZ | *glpK*::*P<sub>J23103</sub>-lacZ-T<sub>rbcL</sub>*; *spe<sup>R</sup>* in WT. |
| WT-J23104-lacZ | *glpK*::*P<sub>J23104</sub>-lacZ-T<sub>rbcL</sub>*; *spe<sup>R</sup>* in WT. |
| WT-J23105-lacZ | *glpK*::*P<sub>J23105</sub>-lacZ-T<sub>rbcL</sub>*; *spe<sup>R</sup>* in WT. |
| WT-J23106-lacZ | *glpK*::*P<sub>J23106</sub>-lacZ-T<sub>rbcL</sub>*; *spe<sup>R</sup>* in WT. |
| WT-J23107-lacZ | *glpK*::*P<sub>J23107</sub>-lacZ-T<sub>rbcL</sub>*; *spe<sup>R</sup>* in WT. |
| WT-J23108-lacZ | *glpK*::*P<sub>J23108</sub>-lacZ-T<sub>rbcL</sub>*; *spe<sup>R</sup>* in WT. |
| WT-J23109-lacZ | *glpK*::*P<sub>J23109</sub>-lacZ-T<sub>rbcL</sub>*; *spe<sup>R</sup>* in WT. |
| WT-J23110-lacZ | *glpK*::*P<sub>J23110</sub>-lacZ-T<sub>rbcL</sub>*; *spe<sup>R</sup>* in WT. |
| WT-J23111-lacZ | *glpK*::*P<sub>J23111</sub>-lacZ-T<sub>rbcL</sub>*; *spe<sup>R</sup>* in WT. |
| WT-J23112-lacZ | *glpK*::*P<sub>J23112</sub>-lacZ-T<sub>rbcL</sub>*; *spe<sup>R</sup>* in WT. |
| WT-J23113-lacZ | *glpK*::*P<sub>J23113</sub>-lacZ-T<sub>rbcL</sub>*; *spe<sup>R</sup>* in WT. |
| WT-J23114-lacZ | *glpK*::*P<sub>J23114</sub>-lacZ-T<sub>rbcL</sub>*; *spe<sup>R</sup>* in WT. |
| WT-J23115-lacZ | *glpK*::*P<sub>J23115</sub>-lacZ-T<sub>rbcL</sub>*; *spe<sup>R</sup>* in WT. |
| WT-J23116-lacZ | *glpK*::*P<sub>J23116</sub>-lacZ-T<sub>rbcL</sub>*; *spe<sup>R</sup>* in WT. |
| WT-J23117-lacZ | *glpK*::*P<sub>J23117</sub>-lacZ-T<sub>rbcL</sub>*; *spe<sup>R</sup>* in WT. |
| WT-J23118-lacZ | *glpK*::*P<sub>J23118</sub>-lacZ-T<sub>rbcL</sub>*; *spe<sup>R</sup>* in WT. |
| WT-J23119-lacZ | *glpK*::*P<sub>J23119</sub>-lacZ-T<sub>rbcL</sub>*; *spe<sup>R</sup>* in WT. |
| WT-lacI-trc-lacZ | *glpK*::*P<sub>lacI<sup>Q</sup>-lacI-P<sub>trc</sub>-lacZ-T<sub>rbcL</sub></sub>*; *km<sup>R</sup>* in WT. |
| WT-trc-theoE*-lacZ | *glpK::P<sub>trc</sub>-theoE\*-lacZ-T<sub>rbcl</sub>; spe<sup>R</sup>* in WT. |
| WT-trc(118)-theoE*-lacZ | *glpK::P<sub>trc</sub>(-10box: J23118)-theoE\*-lacZ-T<sub>rbcl</sub>; spe<sup>R</sup>* in WT. |
| WT-trc(105)-theoE*-lacZ | *glpK::P<sub>trc</sub>(-10box: J23105)-theoE\*-lacZ-T<sub>rbcl</sub>; spe<sup>R</sup>* in WT. |
| WT-trc(114)-theoE*-lacZ | *glpK::P<sub>trc</sub>(-10box: J23114)-theoE\*-lacZ-T<sub>rbcl</sub>; spe<sup>R</sup>* in WT. |
| WT-trc-theoB-lacZ | *glpK::P<sub>trc</sub>-theoB-lacZ-T<sub>rbcl</sub>; spe<sup>R</sup>* in WT. |
| WT-GG | *glpK::P<sub>psbA2</sub>-ggpP*, NSC2::*P<sub>cpc560</sub>-ggpS; spe<sup>R</sup>* and *km<sup>R</sup>* in WT. |
| WT-VBALE-GG | NSC2::*P<sub>cpc560</sub>-ggpS-P<sub>psbA2</sub>-ggpP; km<sup>R</sup>* in WT. |
| *E. coli* | *E. coli* DH5α |

To ensure the proper functioning of the isopropyl β-D-thiogalactoside (IPTG)-inducible promoter and theophylline riboswitch, stock solutions of IPTG (1 M) and theophylline (20 mM) were prepared (Aladdin, Shanghai, China). Theophylline was dissolved in the normal MAD medium and subsequently diluted to the desired working concentration prior to use.

### 2.2 Transformation of PCC 11901

Conjugation of PCC 11901 was performed following our previously established method [20]. Briefly, *E. coli* HB101 carrying pRL443 and pRL623 (Helper), and *E. coli* carrying the target plasmids were cultured. Mid-logarithmic cells were collected and washed to remove antibiotics, followed by their mixed incubation with fresh LB medium for 30 min. Mid-logarithmic culture of PCC 11901 (volume × OD_750 nm_ = 2) was harvested and washed via centrifugation, followed by re-suspension of each transformation in 0.2 mL of MAD medium. The cyanobacteria were subsequently combined with the *E. coli* suspension and incubated for 30 min. Finally, the mixture was applied to a sterile filter film (pore size 0.45 μm) on MAD agar plate. The plate was incubated under an illumination of approximately 100 μmol photons m^-2^ s^-1^ for 24 hours, after which the filter film was transferred to a new MAD agar medium supplemented with the appropriate antibiotics (e.g., 20 μg/mL spectinomycin, 50 μg/mL kanamycin). Transformants were observed after 7-10 days of incubation at an intensity of approximately 100 μmol photons m^-2^ s^-1^.

The natural transformation experiment was conducted based on previous studies with modifications [21]. Culture of PCC 11901 in mid-logarithmic growth phase (volume × OD_750 nm_ = 2) were centrifuged and then re-suspended in 0.2 mL of fresh MAD medium. Approximately 2,000 ng of cyclic DNA was added to the suspension, and the mixture was thoroughly mixed. The incubation of the mixture took place for 12 hours at 37°C in a shaded environment. Subsequently, the mixture was spread onto a filter film on antibiotic-free MAD agar medium. The incubation of the spread mixture occurred at 37°C under a light intensity of 100 μmol photons m^-2^ s^-1^ for 24 h. Then the filter membrane was transferred to MAD agar medium supplemented with the appropriate antibiotics (20 μg/mL spectinomycin, 50 μg/mL kanamycin). The transformed cells became visible within 7-10 days of incubation under the same conditions.

### 2.3 Chlorophyll Fluorescence Induction Kinetics (OJIP) measurement

The protocol proposed by Hendrik et al. [22] was improved to determine OJIP Chlorophyll Fluorescence Induction Kinetics. PCC 11901 was inoculated to OD_750 nm_ = 0.1. After 48 h of cultivation, the culture (3 mL) was collected to place in the dark for 15 min. Subsequently the OJIP parameters were measured using the AquaPen-C (AP 110-C, Drásov, Czech Republic). The data from the wild-type (WT) strain served as the control, and the ratio of each strain to the WT was calculated.

### 2.4 Absorption spectra

Cell samples were collected from PCC 11901 cultures after 48 h of cultivation with an initial OD_750 nm_ of 0.1. The samples were then diluted to an OD_750 nm_ of 0.5 and subsequently normalized. Absorption spectra were measured using a spectrophotometer (UV1601, Rayleigh, Beijing, China). The obtained data were analyzed using UVprobe software (Version X) [23].

### 2.5 *β*-Galactosidase activity measurement

β-Galactosidase (LacZ) activity was assessed based on a previously conducted study [24]. At the mid-logarithmic growth phase of cyanobacteria, samples were collected (volume × OD_750 nm_ = 0.1, or 0.5 for inducible promoters). These samples were then centrifuged and suspended in 1 mL of Z Buffer. To disrupt the cellular structure, 50 μL of 0.1% SDS and 50 μL of chloroform were added and mixed thoroughly. Next, 200 μL of ortho-nitrophenyl-β-D-galactoside (ONPG, 4 g/L) was introduced. The reaction mixture was incubated in a thermostatic reactor (Eppendorf, Hamburg, Germany) at 30°C and 750 rpm for 0.5 min (for inducible promoters, a longer incubation period was necessary). The reaction was halted by adding 500 μL of 1 M Na_2_CO_3_ and thorough mixing. The final solution was centrifugated at 14,000 × *g* for 1 min, and the supernatant was collected to measure the absorbance at OD_420 nm_. Importantly, the corresponding WT result was employed as a blank control. The activity of β-galactosidase was calculated using the Miller value (Miller = 1000 × OD_420 nm_ / (total cell culture × reaction time × cell OD_750 nm_)). It should be noted that the absolute Miller values of LacZ cannot be compared across different experimental batches due to variations in sample losses during centrifugation, the extent of cell fragmentation, and other factors.

### 2.6 GG measurement

Based on the methodology outlined in previous studies with minor modifications [25–27], the PCC 11901 cultures (volume × OD_750 nm_ = 1) was used for analysis. The culture medium was centrifuged at 12,000 × *g* for 5 min to separate the supernatant. The supernatant was then filtered through a 0.22 μm filter to determine the extracellular GG content. The remaining cells from the centrifugation were resuspended in an equal volume of ultrapure water. Subsequently, the cells were subjected to three cycles of frozen-thawed using liquid N_2_ for efficient snap-freezing. Afterward, the samples were once again centrifuged at 12,000 × *g* for 5 min. Transfer the supernatant to a new tube, and the precipitate underwent the same extraction process as described earlier. The resulting supernatant was combined, and the solvent was evaporated using a vacuum concentration system (Her-exi, Hunan, China). Each dried sample was dissolved in 0.2 mL of deionized water and filtered through a 0.22 μm filter for further determination of intracellular GG content. Subsequently, the GG samples were analyzed using an Agilent 1260 series binary HPLC system (Agilent Technologies, CA, USA) equipped with an Aminex HPX-87C column (300 × 7.8 mm, 9 μm; Bio-Rad, CA, USA). The column was maintained at 40°C, and the water flow rate was set at 0.2 mL/min. The same method was employed to analyze various concentrations of GG standards, allowing for the generation of a standard curve. This enabled the quantitative determination of the GG content in the samples.

## 3. Results

### 3.1 Construction and characterization of cobalamin (V_B12_)-independent PCC 11901

Although PCC 11901 exhibited several excellent properties making it one of the most promising cyanobacterial candidates for biotechnological applications, a notable drawback is its reliance on expensive cofactor cobalamin (V_B12_) for optimal growth [18]. Like other marine cyanobacteria, PCC 11901 possesses a V_B12_-dependent methionine synthase (MetH) [28], but the previously isolated strain was found to be an auxotroph. As a result, the initial isolate could not sustain normal growth without V_B12_ supplementation [18]. To address the issue, we replaced the original *metE* gene with a heterogenous one derived from *Synechococcus* sp. PCC 73109 (PCC 73109) [28], which was driven by a constitutive promoter P*_cpc560_* (i.e., the strain WT-metE; **Fig. 2A**). As shown in **Fig. 2B**, growth of WT-metE without V_B12_ supplementation was similar to that of WT with V_B12_, suggesting the successful implementation of our strategy. In addition, we used laboratory-based adaptive evolution to subculture PCC 11901 continuously in medium without V_B12_. Finally, we obtained the strain WT-VBALE, which was able to grow normally under V_B12_-free conditions (**Fig. 2B**). Furthermore, we sequenced the *metE* gene-related region of WT-VBALE. The sequencing results revealed a two-point mutation (TT->AA) 320 bp upstream of the start codon (**Fig. 2C**) and no other mutations were detected in the flanking region. These results are consistent with a recent study of Mills et al. [19] in which we conducted a series of passages of the strain in liquid MAD medium. During the 3^rd^-5^th^ generation of culture, the strain exhibited a notable delay in pre-growth. However, it resumed normal growth after the 6^th^ generation. Finally, as no antibiotic marker existed in WT-VBALE, this strain can be used in place of WT-metE.

**Fig. 2.**
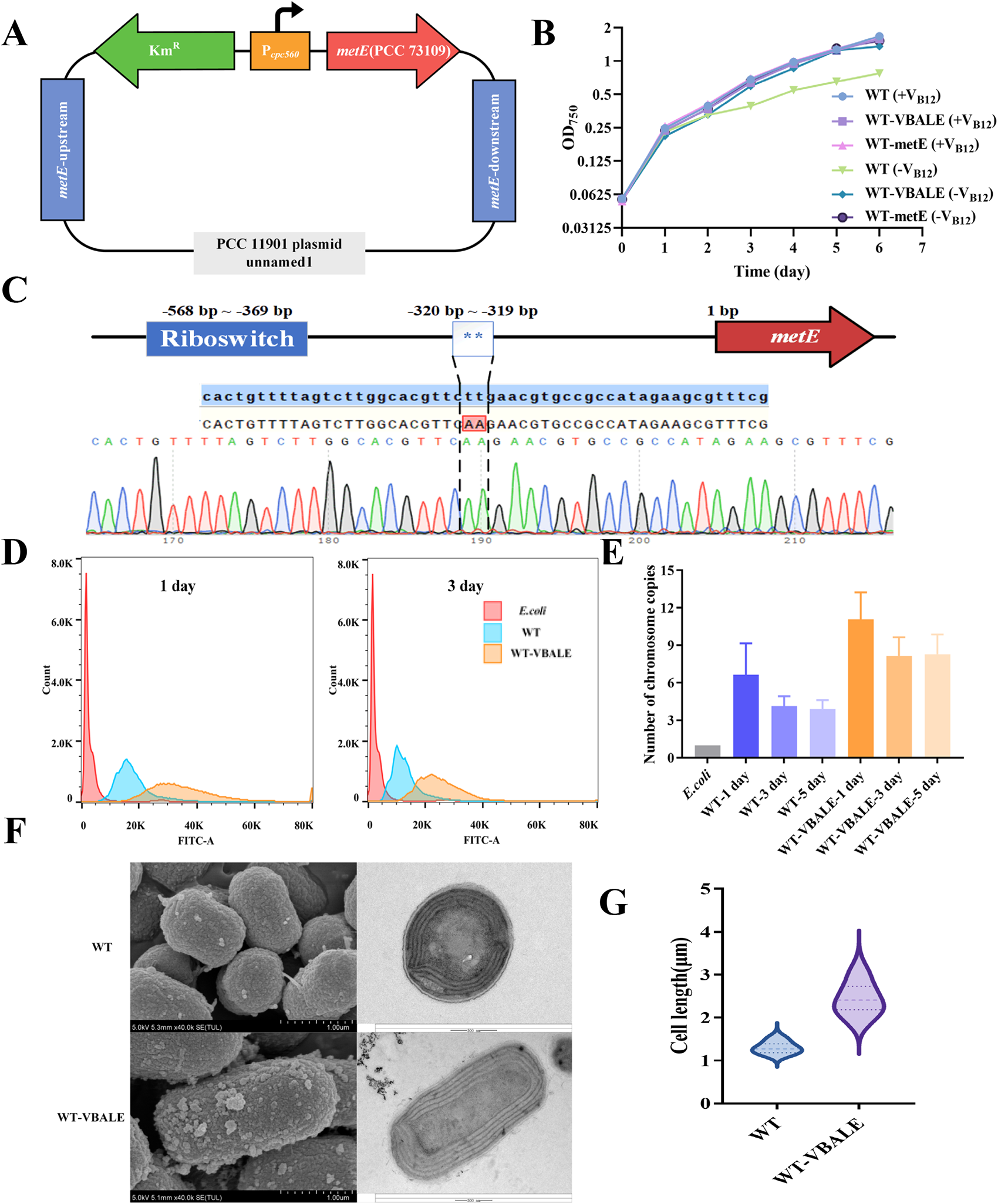
Construction and characterization of cobalamin (V_B12_) independent strains. The error bar represents the standard deviation of the three biological replicates for each sample. **A**) Schematic of the construction of WT-metE; **B**) Growth patterns of WT, WT-metE, WT-VBALE with or without V_B12_, respectively; **C**) Sequencing results of *metE* in WT-VBALE compared with the sequence of WT; **D**) FITC comparison of WT, WT-VBALE and *E. coli* at day 1 and 3; **E**) Relative chromosome copy number comparison of WT, WT-VBALE and *E. coli* at day 1, 3 and 5; **F**) Morphology of WT (top) and WT-VBALE (bottom) cells in the microscope, from left to right, for scanning electron microscopy, and transmission electron microscopy. **G**) Comparison of cell lengths of WT and WT-VBALE.

The presence of a natural genetic transformation system in cyanobacteria is advantageous for genetic manipulation. However, the polyploid of the genome made it difficult to achieve homozygous transformant and aggravated the instability of engineered strain [29]. Polyploid is a common feature for many cyanobacterial species but has not been investigated in PCC 11901. Therefore, we employed flow cytometry to assess the fluorescence signals of WT and WT-VBALE and subsequently calculated their relative chromosome copy numbers. We discovered that the chromosome number was changeable in PCC 11901, e.g., higher in the pre-logarithmic phase (the 1^st^ day) than that in the middle (the 3^rd^ day) or end (the 5^th^ day) of the logarithmic phase for both WT and WT-VBALE (**Fig. 2D**, **2E**, **S1**). Interestingly, the results showed an increased chromosome number in the WT-VBALE mutant compared to WT, as the genome copy numbers varied from 3.90 ± 0.72 to 6.65 ± 2.50 in WT while they changed from 8.14 ± 1.49 to 11.09 ± 2.16 in WT-VBALE, respectively (**Fig. 2E**). The observed trend in chromosome number alterations aligns with earlier findings documented in PCC 7942 [30] and PCC 6803 [31]. The result would be valuable in guiding the screen of homozygous transformants after integrating heterogenous cassettes into the genome of PCC 11901.

As previous studies have demonstrated a correlation between alterations in cell morphology and chromosome numbers [32–34]. Therefore, we examined the morphological changes using both light and electron microscopy for WT and WT-VBALE (**Fig. 2D**, **S2**). It was observed that WT-VBALE exhibited a more elongated shape compared to the WT. We quantified the cell lengths of WT and WT-VBALE based on electron microscopy data (**Fig. 2E**), and the results indicated that the cells of WT-VBALE (average length: 24.44 μm) were significantly longer than those of WT (average length: 13 μm). Additionally, WT-VBALE displayed a rougher cell surface and possessed a higher density of pilus than WT. The differences of cell length and surface between WT and WT-VBALE may influence their self-sedimentation capability, which was explored in following section 3.4 and 3.5.

### 3.2 Transformation optimization and identification of neutral sites

The natural transformation method has been demonstrated to be applicable in PCC 11901 [18, 19]. However, we found it is of low-efficiency as only very limited number of transformants could be obtained typically (**Fig. 3A**). Conjugation, another efficient transformation method to introduce DNA from *E. coli* into cyanobacteria, has been proven effective among several model cyanobacteria [35–38]. In this study, we explored the conjugation in PCC 11901 based on our previous methods developed for UTEX 2973 [20]. As shown in **Fig. 3A** and **3B**, the surface of the resistance-containing medium of conjugation exhibited a higher number of single colonies (mean 54.33 ± 10.07) compared to the surface of the naturally transformed plate medium, which displayed fewer single colonies (mean 17.00 ± 3.61), demonstrating that conjugation exhibited much higher efficiency than natural transformation. The conjugation method was thus employed in all our subsequent studies.

**Fig. 3.**
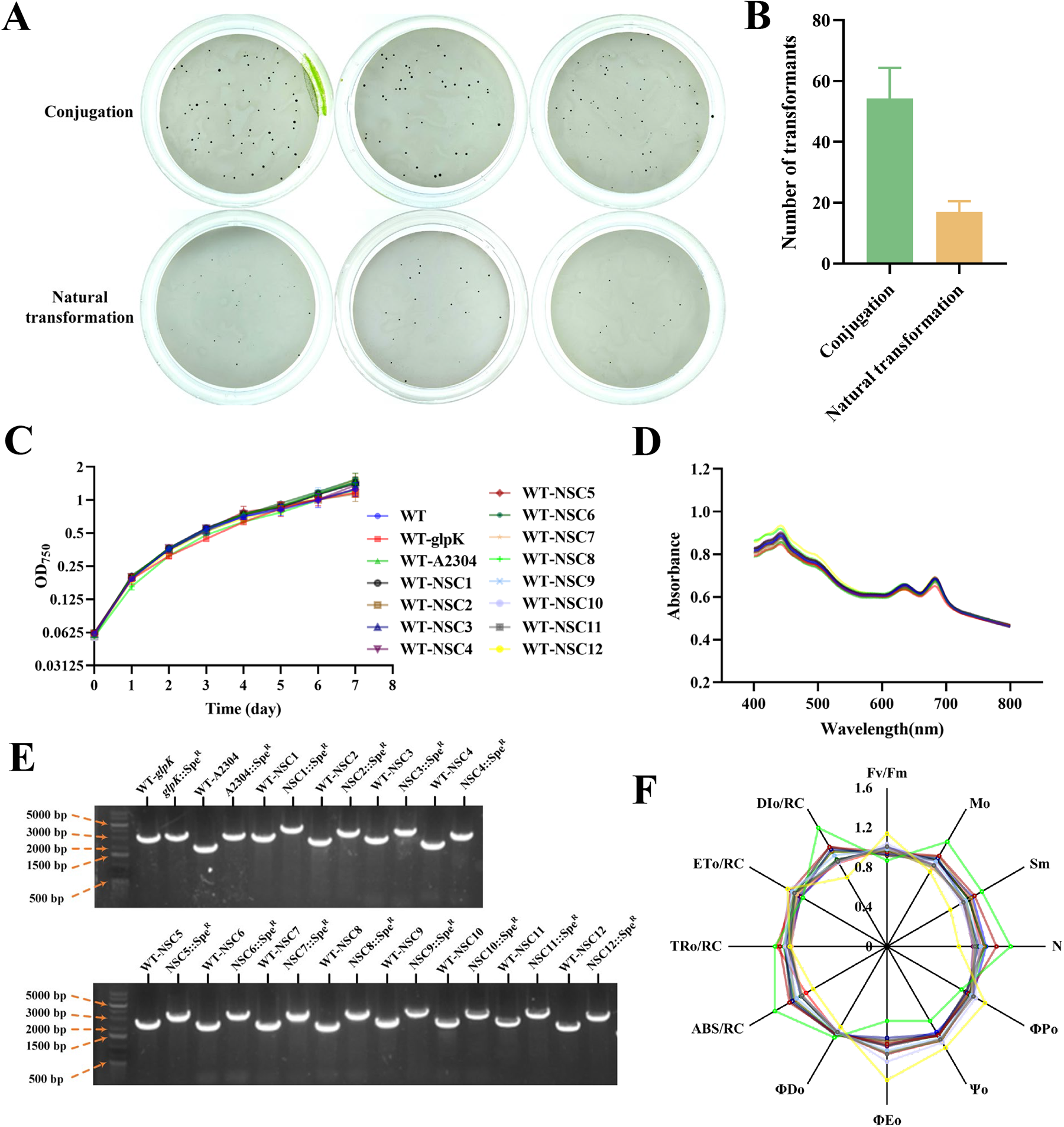
Comparison of transformation methods and characterization of neutral sites. The error bar represents the standard deviation of the three biological replicates for each sample. **A**) Comparison of conjugation (top) and natural transformation (bottom); **B**) Comparison of average transformants between conjugation and natural transformation; **C**) Growth patterns of WT and all neutral sites tested strains; **D**) Absorbance spectrum of WT and all neutral sites tested strains at 48 h; **E**) Agarose gel electrophoresis of all neutral sites tested strains and WT; **F**) OJIP chlorophyll fluorescence kinetic parameters of all neutral sites tested strains and WT at 48 h.

Only three neutral genetic sites were previously reported in PCC 11901 by Włodarczyk et al. [18], including *acsA*, *psbA2* and *fadD*, limiting the sophisticated genetic manipulations. Therefore, expanding the neutral sites of PCC 11901 could be critical for its future commercial applications. Given the high average nucleotide identity (96.76%) between PCC 11901 and *Synechococcus* sp. strain PCC 7002 (PCC 7002) [18], research on identifying neutral sites in PCC 7002 can serve as a reference.

Therefore, glycerol kinase gene *glpK* tested in PCC 7002 was selected in this study [39]. Furthermore, a potential neutral site A2304 in PCC 7002 identified previously by Wang et al. [40] was also investigated. Meanwhile, 12 other intergenic regions (NSC1-NSC12; flanking genes had opposite transcription direction) were chosen as hypothetical neutral sites for testing. **Table S2** presents the sequences of all potential neutral sites. Then a spectinomycin-resistant cassette was respectively integrated into these regions. First, their growth patterns were compared to that of WT and no significant differences were observed (**Fig. 3C**). Encouragingly, we found homozygous transformants could be obtained within just one passage for all these tested neutral sites (**Fig. 3D**). Furthermore, absorption spectra and OJIP chlorophyll fluorescence kinetic parameters of all the mutant strains were also measured. As shown in **Fig. 3E** and **3F**, all the mutants except WT-NCS8 and WT-NCS12 shared the similar characteristics to that of the WT. Together, our results demonstrated that the 10 intergenic regions were suitable to serve as neutral sites for further engineering efforts in PCC 11901.

### 3.3 Characterization of constitutive promoters and inducible promoters

Only two promoters, namely P*_clac143_* and P*_cpt_*, have been previously reported for PCC 11901 [18]. In this study, we evaluated the activity of 22 constitutive promoters in PCC 11901, including all promoters in the P*_J23_*_-_series and the commonly used P*_cpc560_* and P*_psbA2_* promoters in cyanobacteria. A reporter gene *lacZ*, which encodes β-galactosidase, driven respectively by these promoters was intergraded into the neutral site *glpK* (**Fig. 4A**; **Table 1**). The strongest promoter was found to be P*_J23119_*, known for its high gene control expression in other bacteria like *E. coli*, with a Miller value of 16,561.52 ± 4,151.35 (**Fig. 4B**). Conversely, the weakest promoter was P*_J23103_*, with a Miller value of 20.34 ± 2.12, representing a wide expression range of about 800 folds. Notably, the expression levels of the commonly used promoters in cyanobacteria, P*_psbA2_* and P*_cpc560_*, were considerable with Miller values of 5,286.86 ± 692.72 and 4,411.824 ± 575.94, respectively. The β-galactosidase activity controlled by P*_cpc560_* was significantly higher than that demonstrated in the PCC 6803 and the PCC 7942 [20, 24]. This observation might be related to the rapid growth property of PCC 11901, which might possess a higher protein content in cells.

**Fig. 4.**
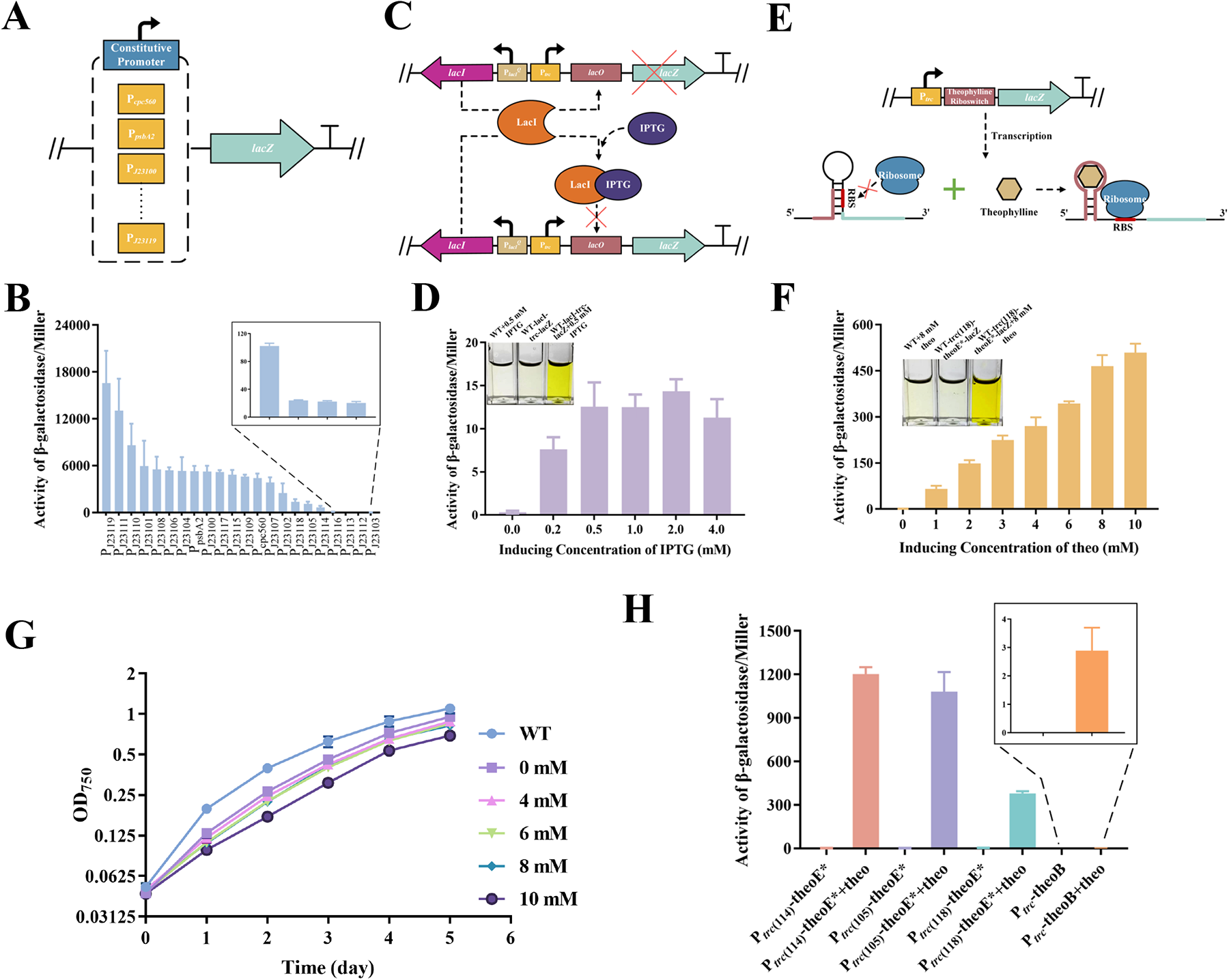
Characterization of various promoters by β-galactosidase activity analysis. The error bar represents the standard deviation of the three biological replicates for each sample. **A**) Schematic of the constitutive promoters; **B**) Activity of β-galactosidase under the control of constitutive promoters at 48 h; **C**) Schematic of IPTG inducible promoter; **D**) Activity of β-galactosidase under the control of IPTG inducible promoter at 48 h supplied with different concentrations of IPTG at 0 h. The cuvette photo showed the results of LacZ activity of WT, WT-lacI-trc-lacZ (without inducer) and WT-lacI-trc-lacZ + IPTG (with inducer) at 48 h; **E**) Schematic of theophylline inducible promoters. These riboswitches regulate downstream gene expression at the translation level by blocking the ribosome binding site (RBS) of mRNA in the absence of theophylline ligand binding. However, when theophylline is present, the RBS becomes available, allowing translation to proceed; **F**) Activity of β-galactosidase under the control of P*_trc_*_(118)_-theoE* at 48 h supplied with different concentrations of theophylline (use theo instead in the figure) at 0 h. The cuvette photo showed the results of LacZ activity of WT, WT-trc(118)-theoE*-lacZ (without inducer) and WT-trc(118)-theoE*-lacZ + theo (with inducer) at 48 h; **G**) Growth patterns of WT and WT-trc(118)-theoE* supplied with different concentrations of theophylline at 0 h; **H**) Activity of β-galactosidase under the control of theophylline inducible promoters at 48 h supplied with 8 mM theophylline at 0 h.

In addition, we evaluated several inducible promoters in PCC 11901. Initially, we created a complete set of isopropyl β-D-thiogalactoside (IPTG)-inducible promoter P*_trc_* based on the Lac operon (**Fig. 4C**; **Table 1**). The strong promoter P*_J23111_* or the medium-strength promoter P*_cpc560_* was used to control the expression of repressor encoding gene *lacI*. Although the construction was successful, β-galactosidase activity was not measured after induction (data not shown). We postulated that the expression of LacI was too strong to be completely bound by IPTG. Consequently, we attempted to utilize weaker promoters like P*_J23118_* and P*_J23114_* to drive the expression of *lacI*, but there was still no β-galactosidase activity detected (data not shown). Instead, we replaced the promoter with a P*_lacI_*-lacI mutant P*_lacI_^Q^* from *E. coli*, which has been successfully used in PCC 6803 and PCC 7942 previously [41–44]. We then constructed the strain WT-lacI-trc-lacZ (**Table 1**). As displayed in **Fig. 4E**, it was shown that β-galactosidase activity remains relatively consistent when the IPTG addition exceeds 0.5 mM. Overall, the expression level of this inducible promoter was low with a minimal amount of leaky expression. Furthermore, we evaluated the theophylline-inducible promoter comprising P*_trc_* coupled with a theophylline-inducible riboswitch, which has been widely used in cyanobacteria previously [20, 45–48] (**Fig. 4D**). We introduced an inducible promoter based on theoE* in PCC 11901 and created the strain WT-trc-theoE*-lacZ. Unfortunately, the element yielded no expected results, displaying significant leaky expression (**Fig. S3**). We hypothesized that the over-expression intensity of P*_trc_*, which has the same -10-box sequence as P*_J23119_* (TATAAT), might cause the failure of the riboswitch. Therefore, we adapted two alternative strategies: *i*) replacing the -10-box of P*_trc_* with the -10-box from the weakly expressed promoters; *ii*) replacing theoE*with a much less leaky B-type riboswitch (theoB) [49]. We evaluated the -10-boxes from P*_J23118_* (-10-box: TATTGT), P*_J23105_* (-10-box: TACTAT) and P*_J23114_* (-10-box: TACAAT) as alternatives by using the last three digits (118, 105, 114) to represent them. We proceeded to construct strains WT-trc(118)-theoE*-lacZ, WT-trc(105)-theoE*-lacZ, WT-trc(114)-theoE*-lacZ, and WT-trc-theoB-lacZ, implementing the aforementioned strategies, respectively (**Table 1**). First, we utilized WT-trc(118)-theoE*-lacZ to determine the optimal concentration for theophylline induction (**Fig. 4F** and **4G**). The results revealed a positive correlation between β-galactosidase activity and theophylline concentration (**Fig. 4F**). However, we observed a slightly negative impact on the growth of this strain when added theophylline exceeded 10 mM (**Fig. 4G**). Based on these findings, we established that the optimal induction concentration of theophylline was 8 mM, a value we employed in subsequent experiments (**Fig. 4H**). Notably, P*_trc_*_(114)_-theoE* showed the highest expression intensity with a Miller value of 1,203.6 ± 47.13. In contrast, P*_trc_*-theoB showed the lowest expression intensity, with a Miller value of 2.89 ± 0.81, representing a significant difference of approximately 400-fold compared to P*_trc_*_(114)_-theoE*. The result suggested that P*_trc_*_(114)_-theoE* was the most effective inducible promoter in PCC 11901. Importantly, all theophylline-induced promoters showed almost no leakage, suggesting the high strictness. These promoters all allowed precise control over gene expression in response to theophylline, providing a powerful tool for manipulating gene expression in PCC 11901, especially in dealing with the possible toxicity from gene expression.

### 3.4 GG synthesis in PCC 11901

Based on the established toolboxes, we aimed to develop cell factories in PCC 11901. GG is a compatible solute that accumulates in various microorganisms and plants, aiding in their adaptation to salt and desiccation stress [50]. The GG synthetic pathway has been elucidated in PCC 6803, involving two enzymes including GG-phosphate synthase (GGPS) and GG-phosphate phosphatase (GGPP) [51]. In this study, WT-GG and WT-VBALE-GG were respectively constructed by introducing the two genes (*ggpS* and *ggpP*) from PCC 6803 into WT and WT-VBALE. Specifically, the expression of *ggpS* was controlled by P*_cpc560_*, while *ggpP* was regulated by P*_psbA2_* (**Fig. 5A** and **5B**; **Table 1**). To evaluate GG production, the constructed strains were cultivated in parallel with WT and WT-VBALE in light shaker incubators. The results indicated that additional pathways caused no change to the growth of both WT-GG and WT-VBALE-GG (**Fig. 5C**). In addition, GG could be naturally produced in WT and WT-VBALE, reaching a highest production of 101.27 ± 4.93 and 136.81 ± 2.84 mg/L in shaking flasks, respectively (**Fig. 5D**). With the exogenous pathways, GG synthesis was significantly enhanced in both WT-GG and WT-VBALE-GG, achieving a highest production of 159.18 ± 4.70 and 200.60 ± 16.41 mg/L in shaking flask, respectively.

**Fig. 5.**
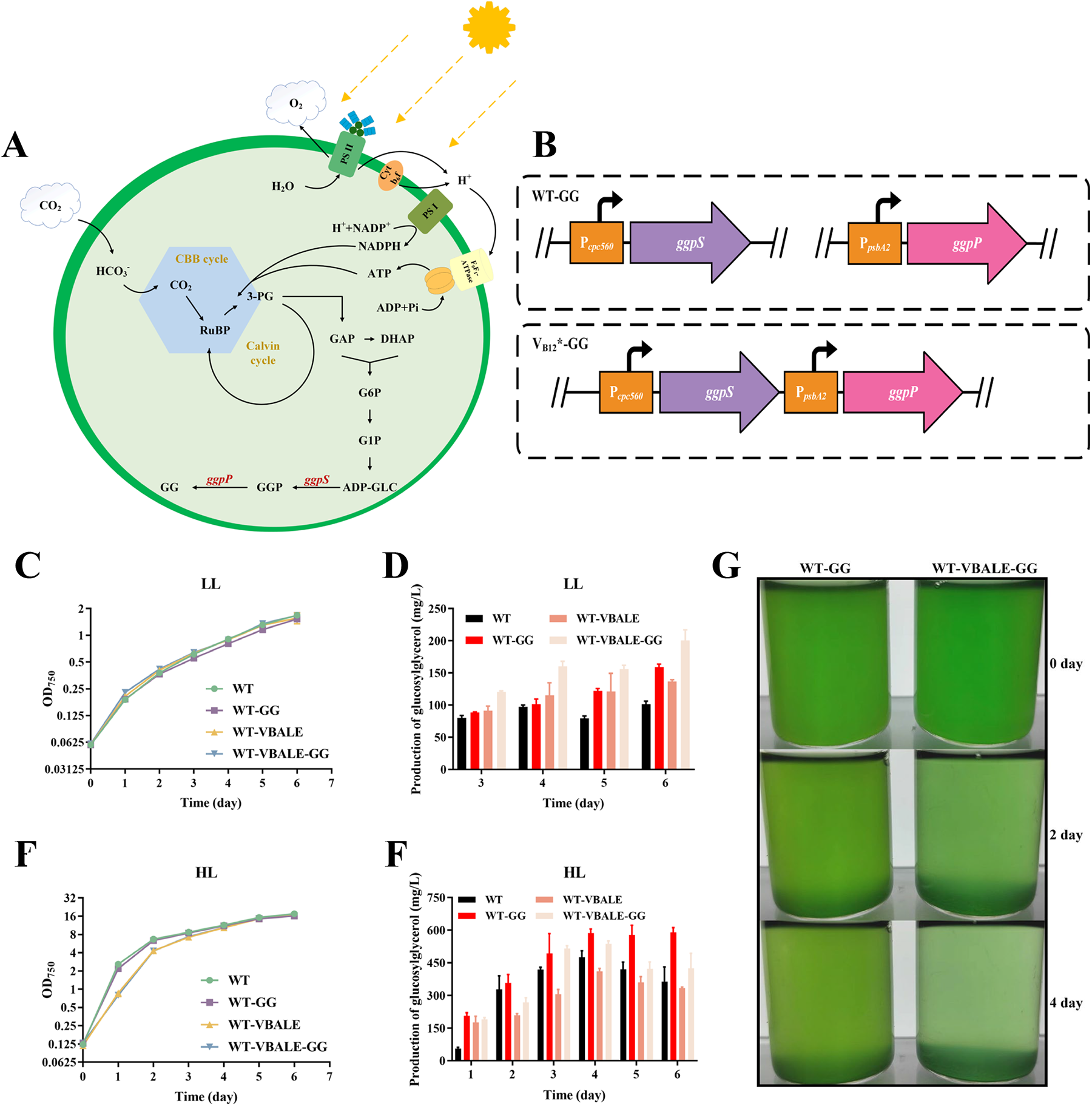
Biosynthesis of GG in PCC 11901. The error bar represents the standard deviation of the three biological replicates for each sample. **A**) Schematic of the GG biosynthesis. Exogenously introduced genes are shown in red. **B**) Schematic of the constructed strains; **C**) Growth patterns of WT, WT-GG, WT-VBALE and WT-VBALE-GG under a light illumination of 100 μmol photons m^−2^ s^−1^; **D**) GG production of WT, WT-GG, WT-VBALE and WT-VBALE-GG at day 3 to 6 under a light illumination of 100 μmol photons m^−2^ s^−1^; **E**) Growth patterns of WT, WT-GG, WT-VBALE and WT-VBALE-GG under a light illumination of 600 μmol photons m^−2^ s^−1^ and supplied with 10% (v/v) CO_2_; **F**) GG production of WT, WT-GG, WT-VBALE and WT-VBALE-GG at day 1 to 6 under a light illumination of 600 μmol photons m^−2^ s^−1^ and supplied with 10% (v/v) CO_2_; **G**) Comparison of self-sedimentation of WT-GG and WT-VBALE-GG at days 0, 2, and 4.

We further investigated the GG synthesis using high-density cultivation in photoreactors with 10% (v/v) CO_2_. As shown in **Fig. 5E**, growth of WT-VBALE and WT-VBALE-GG showed a delay compared to that of WT and WT-GG. The results suggested that though WT-VBALE acclimated to the V_B12_-free medium, they were not totally adapted during high-density culture. Nevertheless, high-density culture significantly improved the production of GG. Detailly, WT-GG exhibited the highest yield of 590.41 ± 21.48 mg/L on the 6^th^ day, while the other three strains (WT, WT-VBALE, WT-VBALE-GG) reached their peaks on the 4^th^ day with yields of 475.81 ± 30.18 mg/L, 410.83 ± 13.69 mg/L, and 537.31 ± 13.37 mg/L, respectively.

In addition, we also investigated the possible utilization of self-sedimentation property found in WT-GG and WT-VBALE-GG given their different morphology described above (**Fig. 5G**). Notably, the sedimentation rate of WT-VBALE-GG was faster compared to that of WT-GG, which is a desirable characteristic for a potential industrial application. It has been reported that thinner and longer cells could exhibit higher sedimentation rates due to the gravity [52], which may potentially explain the superior self-sedimentation observed in WT-VBALE-GG. It was noteworthy that WT-VBALE-GG exhibited superior production using shaking flask and comparable production using photoreactor compared to that of WT-GG. This suggested that the V_B12_-independent mutant strain WT-VBALE held potential as an excellent candidate for industrial applications rather than WT in the future.

## 4. Discussion

As a fast-growing marine cyanobacterium with notable traits such as salt, light and heat tolerance, PCC 11901 exhibits substantial potential as an important chassis for biotechnological applications. However, the need of V_B12_ for large-scale cultures would unavoidably increase the cost, decreasing the utilization feasibility of PCC 11901 for any commercial-scale application. Previously, Pérez et al. [28] mitigated the nutritional deficiencies of PCC 7002 by incorporating the expression of *metE* from PCC 73109, a gene encoding methionine synthase independent of V_B12_. Recently, Mills et al. [19] conducted directed evolution to address PCC 11901’s dependence on V_B12_, obtaining a mutant that could grow in V_B12_-free medium. In this study, we confirmed the feasibilities of both strategies. Furthermore, though growth of WT-VBALE without V_B12_ was comparable to that of WT with V_B12_, we found that these two strains exhibited different morphology. Not only did WT-VBALE became elongated, but its surface displayed increased roughness, which could be due to enhanced extracellular secretions. Additionally, the number of pili in WT-VBALE was increased. Meanwhile, we found the growth of WT-VBALE lagged that of the WT during the first 3 day when cultured in photoreactors. As V_B12_ is also required as a key enzyme of cellular one-carbon (C1) metabolism important for production of the universal methyl donor S-adenosylmethionine (SAM), and for folate cycling necessary for DNA synthesis [53], our results indicated V_B12_ had additional roles besides the activation of *metE* in PCC 11901. Therefore, a more comprehensive investigation of V_B12_-independent mutant strains via comparative transcriptomics and mechanisms elucidation is still warranted in the future.

A diverse promoter library with a broad regulatory range is essential for the industrial application of chassis strains. In this study, we developed an extensive collection of constitutive promoters with an impressive regulatory range of 800 folds for PCC 11901 (**Fig. 4B**). Through screening, we identified super-strong promoters such as P*_J23119_* and P*_J23111_*. Additionally, commonly used promoters in other cyanobacteria model strains like P*_cpc560_* and P*_psbA2_* were found to function well in PCC 11901, suggesting their functional conservation among cyanobacteria. To achieve complete inducible expression, we constructed an IPTG inducible system using P*_trc_* in conjunction with the Lac operon in PCC 11901 [44]. Nevertheless, the system exhibited low leakage expression but also demonstrated low inducible expression intensity even after our optimization (**Fig. 4D**). In the future, the expression of LacI can be constantly explored to find its suitable level. In addition, Topp et al. [54] developed several theophylline-induced riboswitches that can be widely applied to different cyanobacteria strains. Previously, Zhang et al. [55] made significant improvements to citric acid production in PCC 7002 by reducing the citric acid cycling flux using a theophylline-responsive riboswitch known as theoE* which reflected rigorously induced expression. This riboswitch exhibited tightly controlled expression upon the addition of 1 mM theophylline. In addition, the theophylline-inducible riboswitch showed strong inducible expression of about 20 folds upon the addition of 2 mM theophylline compared with that without induction in *Anabaena* sp. PCC 7120, although exhibiting leaky expression [56]. However, the constructed P*_trc_*-theoE* showed severe leaky expression in our study. It might due to the inherent differences of host or the differences of culture temperature which could alter the secondary structure of the riboswitch. Alternatively, changing the original -10-box of the riboswitch led to the identification of several inducible promoters including P*_trc_*_(118)_-theoE*, P*_trc_*_(105)_-theoE*, and P*_trc_*_(114)_-theoE*, in response to theophylline. In comparison to UTEX 2973 with 2 mM theophylline added [20], we observed a 3-fold increase in the intensity of theophylline-induced riboswitch expression in our optimized constructions (as measured by Miller values). This strategy can be further explored in future studies by switching to -10-box sequences with moderate promoter strength or employing other riboswitches with reduced leakage expression [49].

Two significant drawbacks of traditional cyanobacterial cell factories are their slow growth and low productivity. In this case, fast-growing cyanobacteria have exhibited superior feasibilities. For instance, production of sucrose achieved 8 g/L by expressing the sucrose transporter *cscB* in UTEX 2973, which was approximately 3-fold higher than that of PCC 7942 [57]. In addition, Lin et al. achieved a limonene production of 16.4 mg/L in UTEX 2973 [58], which was about 8-fold higher than that in PCC 7002 [59]. In this study, we explored the potential of PCC 11901 as a cell factory via using GG as a proof of concept. Previously, Cui et al. [50] realized 51 mg/g of GG in UTEX 2973, while it was 100 mg/L reported in PCC 6803 [26]. Following the introduction of the GG synthesis pathway into PCC 11901, the yields of WT-GG and WT-VBALE-GG were measured at 101.27 ± 4.93 mg/L and 200.60 ± 16.41 mg/L, respectively. Notably, the yield of WT-VBALE-GG in PCC 11901 was found to be 2 folds greater than that in PCC 6803 [26]. Subsequently, under aerated high-density cultivation conditions, the highest GG yield achieved was 590.41 ± 21.48 mg/L, which was to our best knowledge the highest reported for cyanobacteria. These results highlight the potential of PCC 11901 as a promising chassis for industrial applications, as it demonstrates a relatively high production level through simple genetic engineering modifications. Nevertheless, to further enhance strain productivity, more complex metabolic engineering modifications or the incorporation of different expression control systems should be explored in the near future [60].

## 5. Conclusions

In this study, we obtained a V_B12_-independent strain of PCC 11901 through short-term domestication. In addition, we demonstrated superior efficiency of conjugation compared to natural transformation in PCC 11901. Moreover, we identified 12 neutral sites in PCC 11901 and established the promoter libraries. Finally, we successfully constructed the GG synthetic pathway in PCC 11901 using the genetic toolboxes described above. For future work in PCC 11901, we expect to further expand the genetic toolboxes to include marker-less gene editing methods like CRISPR/Cas and gene regulation utilizing sRNA. Furthermore, optimization of the cell factory constructed in this study can be performed.

## Abbreviations

ADP-GLC: Adenosine diphosphoglucose
G1P: Glucose 1-phosphate
G6P: Glucose 6-phosphate
GAP: Glyceraldehyde 3-phosphate
GG: Glucosylglycerol
GGP: Glucosylglycerol-phosphate
3-PG: 3-Phosphoglyceric acid
RuBP: Ribulose-1,5-bisphosphate
Synechococcus: sp. PCC 11901, PCC 11901
wild type: WT

## Declaration of Competing Interest

The authors declare that they have no known competing financial interests.

## Acknowledgements

This research was supported by grants from the National Key Research and Development Program of China (Grant no. 2019YFA0904600), the National Natural Science Foundation of China (Grant nos. 32270091, 31972931 and 32070083).

## Appendix A. Supplementary data

Supplementary data to this article can be found online.

## Notes

### Competing Interest Statement

The authors have declared no competing interest.

### Summary of Updates

The author list has been updated.

