## Supplemental Tables and figures for "Extended toolboxes enable efficient biosynthesis of valuable chemicals directly from CO_2_ in fast-growing *Synechococcus* sp. PCC 11901"

**Table S1 Synthetic biology elements sequences in this study.**

| Elements | Sequences |
|---|---|
| $P_{cpc560}$ | ACCTGTAGAGAAGAGTCCCTGAATATCAAAATGGTGGGATAAAAAGC TCAAAAAGGAAAGTAGGCTGTGGTTCCCTAGGCAACAGTCTTCCCTA CCCCACTGGAAACTAAAAAAACGAGAAAAGTTCGCACCGAACATCA ATTGCATAATTTTAGCCCTAAAACATAAGCTGAACGAAACTGGTTGTC TTCCCTTCCCAATCCAGGACAATCTGAGAATCCCCTGCAACATTACTTA ACAAAAAAGCAGGAATAAAATTAACAAGATGTAACAGACATAAGTCC CATCACCGTTGTATAAAGTTAACTGTGGGATTGCAAAAGCATTCAAGC CTAGGCGCTGAGCTGTTTGAGCATCCCGGTGGCCCTTGTCGCTGCCTC CGTGTTTCTCCCTGGATTTATTTAGGTAATATCTCTCATAAATCCCCGGG TAGTTAACGAAAGTTAATGGAGATCAGTAACAATAACTCTAGGGTCAT TACTTTGGACTCCCTCAGTTTATCCGGGGGAATTGTGTTTAAGAAAATC CCAACTCATAAAGTCAAGTAGGAGATTAATTCA |
| $P_{psbA2}$ | GGTATATGGATCATAATTGTATGCCCGACTATTGCTTAAACTGACTGAC CACTGACCTTAAGAGTAATGGCGTGCAAGGCCCAGTGATCAATTTCAT TATTTTTCATTATTTCATCTCCATTGTCCCTGAAAATCAGTTGTGTCGCC CCTCTACACAGCCCAGAACTATGGTAAAGGCGCACGAAAAACCGCCA GGTAAACTCTTCTCAACCCCCAAAACGCCCTCTGTTTACCCATGGAAA AAACGACAATTACAAGAAAGTAAAACTTATGTCATCTATAAGCTTCGT GTATATTAACTTCCTGTTACAAAGCTTTACAAAACTCTCATTAATCCTTT AGACTAAGTTTAGTCAGTTCCAATCTGAACATCGACAAATACATAAGG AATTATAACCAA |
| $P_{J23100}$ | TTGACGGCTAGCTCAGTCCTAGGTACAGTGCTAGCTATTGTGAGCGGA TAACAATTTCACACATACTAGAGAAAGAGGAGAAATACTA |
| $P_{J23101}$ | TTTACAGCTAGCTCAGTCCTAGGTATTATGCTAGCTATTGTGAGCGGAT AACAATTTCACACATACTAGAGAAAGAGGAGAAATACTA |
| $P_{J23102}$ | TTGACAGCTAGCTCAGTCCTAGGTACTGTGCTAGCTATTGTGAGCGGA TAACAATTTCACACATACTAGAGAAAGAGGAGAAATACTA |
| $P_{J23103}$ | CTGATAGCTAGCTCAGTCCTAGGGATTATGCTAGCTATTGTGAGCGGAT AACAATTTCACACATACTAGAGAAAGAGGAGAAATACTA |
| $P_{J23104}$ | TTGACAGCTAGCTCAGTCCTAGGTATTGTGCTAGCTATTGTGAGCGGAT AACAATTTCACACATACTAGAGAAAGAGGAGAAATACTA |
| $P_{J23105}$ | TTTACGGCTAGCTCAGTCCTAGGTACTATGCTAGCTATTGTGAGCGGAT AACAATTTCACACATACTAGAGAAAGAGGAGAAATACTA |
| $P_{J23106}$ | TTTACGGCTAGCTCAGTCCTAGGTATAGTGCTAGCTATTGTGAGCGGAT AACAATTTCACACATACTAGAGAAAGAGGAGAAATACTA |
| $P_{J23107}$ | TTTACGGCTAGCTCAGCCCTAGGTATTATGCTAGCTATTGTGAGCGGAT AACAATTTCACACATACTAGAGAAAGAGGAGAAATACTA |
| $P_{J23108}$ | CTGACAGCTAGCTCAGTCCTAGGTATAATGCTAGCTATTGTGAGCGGAT AACAATTTCACACATACTAGAGAAAGAGGAGAAATACTA |
| $P_{J23109}$ | TTTACAGCTAGCTCAGTCCTAGGGACTGTGCTAGCTATTGTGAGCGGA TAACAATTTCACACATACTAGAGAAAGAGGAGAAATACTA |
| $P_{J23110}$ | TTTACGGCTAGCTCAGTCCTAGGTACAATGCTAGCTATTGTGAGCGGAT |

| | |
|---|---|
| | AACAATTTCACACATACTAGAGAAAGAGGAGAAATACTA |
| $P_{J23111}$ | TTGACGGCTAGCTCAGTCCTAGGTATAGTGCTAGCTATTGTGAGCGGAT<br>AACAATTTCACACATACTAGAGAAAGAGGAGAAATACTA |
| $P_{J23112}$ | CTGATAGCTAGCTCAGTCCTAGGGATTATGCTAGCTATTGTGAGCGGAT<br>AACAATTTCACACATACTAGAGAAAGAGGAGAAATACTA |
| $P_{J23113}$ | CTGATGGCTAGCTCAGTCCTAGGGATTATGCTAGCTATTGTGAGCGGAT<br>AACAATTTCACACATACTAGAGAAAGAGGAGAAATACTA |
| $P_{J23114}$ | TTTATGGCTAGCTCAGTCCTAGGTACAATGCTAGCTATTGTGAGCGGAT<br>AACAATTTCACACATACTAGAGAAAGAGGAGAAATACTA |
| $P_{J23115}$ | TTTATAGCTAGCTCAGCCCTTGGTACAATGCTAGCTATTGTGAGCGGAT<br>AACAATTTCACACATACTAGAGAAAGAGGAGAAATACTA |
| $P_{J23116}$ | TTGACAGCTAGCTCAGTCCTAGGGACTATGCTAGCTATTGTGAGCGGA<br>TAACAATTTCACACATACTAGAGAAAGAGGAGAAATACTA |
| $P_{J23117}$ | TTGACAGCTAGCTCAGTCCTAGGGATTGTGCTAGCTATTGTGAGCGGA<br>TAACAATTTCACACATACTAGAGAAAGAGGAGAAATACTA |
| $P_{J23118}$ | TTGACGGCTAGCTCAGTCCTAGGTATTGTGCTAGCTATTGTGAGCGGAT<br>AACAATTTCACACATACTAGAGAAAGAGGAGAAATACTA |
| $P_{J23119}$ | TTGACAGCTAGCTCAGTCCTAGGTATAATGCTAGCTATTGTGAGCGGAT<br>AACAATTTCACACATACTAGAGAAAGAGGAGAAATACTA |
| $P_{lacI^Q}$ | GACACCATCGAATGGTGCAAAACCTTTCGCGGTATGGCATGATAGCGC<br>CCGGAAGAGAGTCAATTCAGGGTGGTGAAT |
| $P_{trc}$ | ATGAGCTGTTGACAATTAATCATCCGGCTCGTATAATGTGTGGAATTGT<br>GAGCGGATAACAATTTCATAC |
| $P_{trc}$(-10box:<br>J23118) | ATGAGCTGTTGACAATTAATCATCCGGCTCGTATTGTGTGTGGAATTGT<br>GAGCGGATAACAATTTCATAC |
| $P_{trc}$(-10box:<br>J23105) | ATGAGCTGTTGACAATTAATCATCCGGCTCGTACTATGTGTGGAATTGT<br>GAGCGGATAACAATTTCATAC |
| $P_{trc}$(-10box:<br>J23114) | ATGAGCTGTTGACAATTAATCATCCGGCTCGTACAATGTGTGGAATTGT<br>GAGCGGATAACAATTTCATAC |
| theoE* | GGTACCGGTGATACCAGCATCGTCTTGATGCCCTTGGCAGCACCCTGC<br>TAAGGAGGCAACAAG |
| theoB | GGTACCGGTGATACCAGCATCGTCTTGATGCCCTTGGCAGCACCCTGA<br>GAAGGGGCAACAAG |
| *lacO* | TTGTGAGCGGATAACAA |
| $T_{rbcL}$ | ACCGGTGTTTGGATTGTCGGAGTTGTACTCGTCCGTTAAGGATGAACA<br>GTTCTTCGGGGTTGAGTCTGCTAACTAATTAGCCATTAACAGCGGCTTA<br>ACTAACAGTTAGTCATTGGCAATTGTCAAAAAATTGTTAATCAGCCAA<br>AACCCACTGCTTACTGATGTTCAACTTCGACAGC |

**Table S2 Neutral sites sequences in this study.**

| Neutral sites | locus_tag | Sequences |
|---|---|---|
| *glpK* | FEK30_03865 | TTAGGCCCAGTTTTTCGCTCGTTCAACGGCTTTTTGCCAG<br>ACTTTGAAATGGGCTTGGGCTTGGGAGGCGTTGGCACTG<br>GGTTTAAAAACGTAATCAATCTTGCGGTTTTGCACCAGGG<br>TTTGGTAATCATCCCAAAAGCCAACGGCCAATCCGGCGGC<br>AAAGGCGGCTCCCTGGGCCGTGGCATCCAGCACCGCGGG<br>ACGTTCCACGGGAATGCCCAATACATCGGCTTGGAATTGC<br>ATCAGAAAGTCGTTATTGCACGCGCCGCCATCGACTTTTA<br>ATTCTTGGATCGGGGTACCAGAATCTTGGTTAATTGCTTCG<br>ACTACTTCTTTGGCTTGGTAGGCGATCGCCTCTAGGACAG<br>CCCGCACCATGTGTTCTTTTTTGACGCCTCTCGTTAAACC<br>GAGAAATGCGCCCCTGGCACTCATATCCCAGTGGGGCGCT<br>CCCAGGCCACTCAGGGCAGGGACAAAATAAGCGCCGCCG<br>TTATCGTTCACCCCTTGGGCAAGATCATTCGTTTCCGCCGC<br>CGTTTCGATAATTTGTAGGCCATCGCGCAACCATTGGATAC<br>AAGCCCCAGCGGTGAACATACTCCCTTCGAGGGCATAGCC<br>AATATCGAGATTTGTGCCATCGTGATGGGCTTGGGTGCAG<br>GCGATGGTCGTTAGGAGCTTATGCTTGGAGCGTTTAATCT<br>CGTCGCCCGTGTGCGCCACCAAAAACGCTCCTGTGCCATA<br>GGTGCATTTCAATAAGCCGGGGCGATCGCAACCATGGGCA<br>TAGAGGGCCGCCTGTTGATCCCCAAAGATGGCGGTGATCG<br>GAATTGCCGCCCCCAATAAACTTGGATCGGTTTTGCCAAA<br>TTCCCCTAGGCTCGACTGCACCGTTGGCATCATCTGGCGG<br>GGAATATCGAATAAATCCAACAAATCCGGATCCCAATCTTT<br>TTGGTTGAGATTGAGCAACATTGTCCGGCTGGCGTTGCTG<br>TGATCTGTGGCGTGGACTTTGCCTCCCGTGAGGTTCCACA<br>AGGCCCAGGTGTCAATGGTTCCAGCGAGGACATTATTGGG<br>GTTAATGGCACTATTTTCCTTAGCCCAGCTTAATAACCAAT<br>TCAGTTTCGTTGCCGAAAAATAAGCATCCAACACCAAGCC<br>CGTTTTATCGTAAATTTCTGCCGCTTTGCCTGCCGCACTGA<br>GGGTTTGACAGAGGGGAGCCGTCCGCCGATCCTGCCAGA<br>CAATTGCTTTATGAAGTGGTTTGCCAGTGGTTTTATCCCAG<br>AGGACACAGGTTTCCCGCTGCACCGTCAGGCCGATCGCC<br>GCGATATCCTGGGTTTCAATGCTGCTATTGTTGACGACCTG<br>TTGCATGACGGCTTTGGTGTCGTTCCAAATTTCCGTCGCAT<br>CATGTTCGACCCAACCGGGCTGGGGATAATATTGGGTTAA<br>TTCTTTATAGGCTTGGGCAACAATATCGCCTTTGTGGTTAA<br>ACAAAAGGGCACGATTACCAGTGGTGCCGAGGTCTAACG<br>CAAGAATGTATTTTTGATGGGCCAT |
| A2304 | FEK30_15790<br>to<br>FEK30_15795 | TTAATTGCGGTCAATTTTTAAAAAACTTTTAAAAAAAATC<br>GGCTGCCAACGGCCTGAATACTCCTGCCATTCCAAGGAAT<br>ATCTAGATCAGAAAAAATCTCCCCAGTCACTTTCGCCAAC |

| | | |
|---|---|---|
| | | AAAAAACCAGAGCTTGCTGACTCTGGCAC |
| 09285 | FEK30_09285 | ATGTATCAAGCTGTTTATACTAAGCGTTTTGAAAAGGACAT<br>TAAAAAAGTCAAAAAGAGGGAAAAGATTTAGAAAAGTTC<br>AAGCAAGTTGCTAGGGTATTAATTTTAGGTCAAACTTTAG<br>AGGCAAAGCATCGAAATCATAAATTGATTGTCAATTATCA<br>AAATCGTCGAGAATGTCATATTGAGCCTGATTGGTTATTGA<br>TCTATAAAATTGAACAAGAGCAAATTGTTTTTGAGCGAAC<br>AGGTAGTTATTCAGACTTATTTGATTAG |
| 09935 | FEK30_09935 | TTACTTTAAACCAGCTTGTTTGAGAATACTATTTTGGGTTC<br>CAGGAGCTAGTTCGTCACTGAATTTTCCTGCAACTGTTAC<br>TCATCCTAGTTTAGTTGGATGCCTGTACTGACGGTGACTTC<br>CGCGAGTTCTAGTGAGAATCCAACCATCTTGTTCTATGAG<br>TTTGATAATTTCTTTAACTTTCAT |
| NSC1 | FEK30_01740 to FEK30_01745 | ACCGTTGAGCTCGTTCCCCTTCCTTTAGGGACAATCAACA<br>TACAAGTCAGCGATCGCCGCTGATTTTTTTGCCCATTTTC<br>AGAAGACTTGTACCCCAAAAAAGCCCTTGAATCTTTGAC<br>GGCGGCTACATATAATATAGAAAAGATAGTTTCTGTATCAA<br>AAGTAACTAAATCAAGTTTTTCTTAAAGGGAGTTTAAGGC<br>TATGCGGATGCGTAAAGAAAAAAATGAAACCCTCCGGGC<br>GATGGAACGGGTTTGGGCGAGACATATGCTCAAGCAGCA<br>AGATGCCCAGAACAAAAATGCGGATGCAGTCCGGGAAGC<br>AAAGGAAATTCTTAACGAATACGGCGAATAAGTCAGTTAA<br>ATAATTTTTACGGTAGGCATATGTGTTCAAACATATGCTTTT<br>TTCATGACCTGAAACTCCAGAATGGG |
| NSC2 | FEK30_02620 to FEK30_02625 | TCGAGGCCGCAAAAAGTGTAGTAACCGCCGTCATTGCAC<br>CTTTTCTGGTGGGTGGGGTGCTGTTGGGGACTTGGACATA<br>TTTGACGAAAACTGGTGGCGATCGCTTAATGAGTACCACA<br>AAATAGGTTTAGCCTTGTTTTAAGACGAATCTTAACCCTG<br>GATCGGCTCAGCCAAAGACATCTGCTGCCAAAGGCTGAC<br>AATTAAGTTTTGTAATCCTGAGTGCGTCATCTGTCCAAAG<br>GCAGATCGCTAGATAGACATCGTTAAGAAATGCAGATATG<br>ATGAAAGAACAGTAAAGAAAAACCATCTTTTTGTAATGGA<br>AAAATAAAGTACATATACTTACTGGCAAAAGTAGCTCTTA<br>AAAAAAGGAGATCGCCGCTAGAGCTTGTCGTTAACAAAA<br>ACATTTTTTCTGCGCTGTTATCGGCCAGTTGTTGTGCCAAC<br>TGACCCAACAAGATTCAAAAAACCAAATAAGACAAAGTT<br>TATTTTTTTAGAGAACTTCTTTAAAAGAAACTGACTTAGG<br>GGAACCTCGATTAATTGCTTTTTTGACGATTTTTTGAGCCT<br>GAAACAACGAGGATACGTTGCCGATCTGGCGAAATTTTG<br>AATTTTTCGAGGTTCCCTTAGGTGTCTTTTTTTGATCAAGG<br>AGATGCCCTCTTTGTGGTAAAAAAATCAAAAGTAACCCAC<br>AGATTGCTCGATTTTTACCGACGATTTTGTGTGCAAATACA<br>AAATATTTAATCGATTTAACTTGATATCCGTAATATCAATTA<br>GGTATTCTTGCTTGGTACTGTTTTGACCGTAAAAATACCTA |

| | | |
|---|---|---|
| | | AAAATGCCCTTAAAACCACTGTACTTAGGTGCAATTAATAT<br>CTTTTAAAACATATAAATCAGTGAAAATAACTGCTTTTTCT<br>TGTGTTTTTACCTGAAAACAATGGACTTTATGGACTTTTGT<br>ATTTTTGTTTTTTTTGCTATTAGGTTTTGGATCTCTTTTGGG<br>TATGATTAGGAGTATAAAGCAAAAAGAAATTTGTCAAGCC<br>AGACGATCACTGCCTCAAATTCAAAAATGCTTTACAGGTC<br>AGAAGTTTTTTGTTTTAAAACTGAGGAAAAGCC |
| NSC3 | FEK30_02965<br>to<br>FEK30_02970 | TATCAAAAGACCTCAAGGATTAGATAAAAGGATTCAGATA<br>GGGACAAGAGTATACAGTGAAATCAAACCCAGAATGATA<br>ATCATAATCATTTTTGTTTGATTATCAGTTCACTATAAGCAA<br>ACTAAGAGGAGATAAACTTCTGTGGAGAAAGTAGACTGG<br>CGGCGATCGCCATACTTGTAGAAATGAGAATATCAAGGTG<br>ATAAGAGCAGACTGACACGCTACTCCCTTGGGATATACCG<br>GAATAAAAAGAGATGAGTATGCTGTGCAAGTTAAGAAAC<br>AGTGTCGATCAAGCTTTCATCTCAATTGGGGATCGCATCCT<br>TCTACAACAATGGTTTATCTAACGCAAACACGGGCAGGCT<br>GAGTGTGTTCGCCGCTGGGTACGTCCCATGAGAGACCTTA<br>AACTCGCTGCAGCGAATCCTTGGGTTGAGTGCATTACAAA<br>ATTCTCCACACACTAAATTTACGAAAACAATAATGATAATC<br>GTTTTCGTAAAAGTCTACAAGAGAGATCGTCTTTTGTGGA<br>GTCTAGTTGGCAAGAATGCTAGCGGGGTGATTCCGCATCA<br>ACTGGCGAAAAACCGAGGCAAGCCGCAAACATAAAGTAA<br>AAGAAATCAATCATTTTTTTATACCGAGAAAATAAAGTTGT<br>TTTAACGAAATATCTGGTCTAACCTCAGGACTCCGCTAAA<br>GCATGGTCTAAACTGAGGGTCGCTAAATAATATTGGGCAA<br>GATCAGGAGCCATATGGGGCGGTCTCCCCTGGGATTGCTC<br>TGGGGCTTGGCTATCTTGGCGGAGCTGCTGGGTAATATCG<br>GTCGTACCGCAGGAGTGGACAATTTTCATGGGATATTTTA<br>ATGAAAATAATTCTTATTTAAACTAGATAAAGTGTATCGAT<br>ACTTGGACAGCTCGTCAATAGTATTGATACTTATTTTCAAT<br>AAAATCCTGGGTGAGTTTGT |
| NSC4 | FEK30_03485<br>to<br>FEK30_03490 | CAAAAGTATTTTCCGCTTTATAATTAAGGGCTATACTGCAC<br>CTGCGACATAACAACTCAAATCTACCAGAATATTACCGCA<br>GATTATAGCTACAGAATAGCGGTCAGGAGACAAATTACAG<br>TTATTTACCAATTGATCACCACTGCCTTTTCCATTTTCTTTA<br>GTGCCACTGAGGTATTTTCAGGCTCAAGCATTGTTTGTCA<br>GGCAGAACGGCCCAAGCCTTTTCCTTCCATGGCCTGAATC<br>AGTTATCAAGCCAATACCCGCTTAAGGGGTAAATCGTTAA<br>CCTCCAAGAAAAAACCGGTATGCTGATTGTTTTTCATTTT<br>GGCGATTTTTGGCGATCGCCTTACGGACATAAATATCGGC<br>ATCAAAAATGGAGAGCTGATCGCGTTGGGTCAAGTAAGT<br>CTTGCTATACTTTTTCCCGTCGCAATGCCCCCCATCAAACC<br>GATGATTCCCGGCTGCGTCACCGTACGCGGATCAGCTTAT<br>AGTTTGGCATGGCTGCCATCACAAAACGGCGCATCCTGGG |

| | | |
|---|---|---|
| | | TGTGCTTACAAAGACACAATGTCACTTTTTTCTTCTCTTCT<br>AGGGTGAACTTTTGGGGCTTGAATTCCGTGCCTTTGTGGC<br>TGCCATCACAAAAGGGTTGATCCGCAGATTTACCACAGGC<br>ACACCAGAAATAATCGCCGGCCTCAAGTTCTAGCGTCATG<br>GGCTTTTTCGCCGCAATTGTTGGTTTAGCCATATTCATTTTT<br>CCTCTATAAACTTTGGGCAATAAACTTGACTGACCTAGGG<br>CAAATCAAACAATAGAAATTCCGTCTCGGCCCCTTGACCT<br>GCAAAAATGAGTTCTGTTTCGCCACTGATGCCAGCCCCAT<br>CGCCCGCTTCCATCGTAACGCCATTCACCGTGAGGCTTCC<br>TTTGACAATCTGTAGCCAAGCC |
| NSC5 | FEK30_06605<br>to<br>FEK30_06610 | GGATTTGGAGACCAAATCCCTACGGGAAAAACGGGACTG<br>GCCGGATTCGAACCGGCGACCTAGCGCTTCAGATTGGTGT<br>GGGTTTCCCCGACTCTCTGGACTATCTCTTCACCCTAGAA<br>CCCAGGATTTGGTAACCAAATCCTACAGTTTCCGTAGGTG<br>GTGGCCGTTATGCTGGGTTGGCGATGACCTTTCGCTACAA<br>ACGTTAGCATTCAAGATCCTCAAAAAAAATTTGCCCCAAA<br>ACCAACCAGCTAGTCTCTGCACCTTCCTCCGGCGAGGCA<br>CCGAAAGGCTTGGCTCAGGGTTACCCTATGATGGAAATTT<br>TCATCACTTAGGCTTCCTTGAATTTGACCACATTCAGCCAT<br>GGCGTTTCCGATCCATGCTGCCCGAAAGATTCAGGAGGCG<br>CTTGCTCTATCCTGCTGAGCCACAGCCCCAATAAAGCGCC<br>AAAATTATAACATAGCTTTTTTTCTATG |
| NSC6 | FEK30_07090<br>to<br>FEK30_07095 | TCCTGAACCCTTATTCCTTCAGAAGAGTTGCCAACATCGC<br>CCGCGAGATCGATAAGCAAATATTACAAGATATAAAGTATC<br>CATCACCGCTACCTATCTTTGCTGATACCTTATAAGCAAGG<br>TACAAAATAAAAGTTCTGAAAAAGCTTTTATTCGGACGAA<br>GAGATTGAGCGGTTGTACCTCCTTTGTTCGGTCTCGCCCA<br>GAAGTCCTTCTCAACATTTCAAGTTGAAGATAGACTCGGT<br>TATTCCCCCTAACGGAATCGGGTAATCGGGCTATAATCACT<br>GCCTGTTAGGTATCGTTTTTCGAGAAGCGCTTGTTGGGTT<br>CGTTGTTAACCTCGTTTTGTTGTTTGGTAAATCTTTTTGTT<br>AGTTAAATTTGGATTGTAAGCCTGCTCCCCTGAGTAGGCT<br>TTCGCCTTGTTTGGGGGAAATCATCCCAAACAGCGGTAGA |
| NSC7 | FEK30_09325<br>to<br>FEK30_09330 | ATTTTGAAATTAATGCCCAAAAAATCAAACTCTGTTTGCG<br>GAGATCGGAAAAAGGTTATCTGCGTTTTTGATTGACTTTT<br>CCAGTCTAATTTGTGAAGGTGTCGCTTTATTGAAGAGCCA<br>GAATATTTGATGATTTCAAGGTTTTGTAGTCTTTGTTTATG<br>GGCATAATCTTTTCACAAAAGAGTCAAAATGACGGCTTTT<br>TCCTGATCCGATTCGGGGTCAATTTCCTTGATTGCCTTTCT<br>GCCGCTACAAAGTCAAAAAAAATCAAGTCGAACCAATTA<br>AAAAAGCACTGAGACTATTCCCAGTGCTGTTGTTTTATTA<br>GATTATTAAAACCAAAATCCCAGTGATGTTTTATTTAAAC |
| NSC8 | FEK30_11495<br>to | ACCCTGTTGCAACTAGCCGCTGGGGTGTATTGATTTAAGA<br>AAACTAAAGACTGTTGTGAATCCTGGGGTGATTGTTCGTA |

| | | |
|---|---|---|
| | FEK30_11500 | GCGATGTAACCCTGGGATATTTTGTGTTGGGCACAGAAA<br>AAAGACCGGCGACTTTCCGTGTTTCGTCTCGGTCTTCTTT<br>CTGGTTCGCTCCTCACACACCTGTAAGTGTAGACCATCTT<br>TTCTCAAACTGCCATGTTTTTTTCAAGTTTTGTGTCGAAA<br>ATTTTAAATAGACCAATTAAAAAAAGCGTGAAAGCTAACG<br>TCCCACTGCCTTTGAGTCTTATAAA |
| NSC9 | FEK30_13325<br>to<br>FEK30_13330 | GCTAATTTAAAATCATGAGCGTTTGTTGGGTGAGGGCGAT<br>CAAGGCGATCGCCCATTGTTTTAAGTAAGGCAATGATTTT<br>CAACGAAATCACGAAAGAAAGCTTTGAGAGTTACGTGGA<br>AGCCGCCGCATCAGTAGGAGGCACATATCGAGGGTTGTTA<br>GCGGCTCATCGAAGGGTAAACTTTGCTGGCCGGTTCAAAT<br>ATCGTGGACAAAAGTGGAGCGCTGGCAAAACTTCTATTTT<br>CCCAATCAAAAAGCATAAAACCCCCTCTAGGCTTTCTGTC<br>CCCCCAGTCGAAAAGCCCACAAATCCGCCTAACCCCTGT<br>CGGGTTCCAAGGCAGAAAAAGGTCTCAGCCAAGCAAATT<br>ACGCAGAGAAAAACCTTGCGGGAATTGCTTAACCTTTAA<br>CTAAAGCGATCGCCTTT |
| NSC10 | FEK30_14635<br>to<br>FEK30_14640 | CGTAAAAAACAAAATAATTCTAATTTAAAGAACTAATTTG<br>TTTGTCTCCCAAATGAATTTATAAGTATTTTTAAAATTGCTT<br>TTGATGGCATTTTCACATAACCATCCAACTCAAACAATGG<br>TGGCTTTTGGGGCGATTTGAAAAGCAAAAGAAGCCTCTT<br>TTAATGATGTCCTACTTTTTTGCCTGGTTTATCGGTTTTTGG<br>TTCATTCTCTTTGGGGGAATGCTCCTTAAAGCAACTATTGG<br>TCAGCAACTACGTCCTTTACTGAGCTACCGTAAGCCTCCC<br>CTGGGCCAAATTCGCCCTGGTGAGAAAATTTATTTTCAGG<br>GACAATATAGCGTTGGCAAAAGTATTCTTTCTCCCTTAACA<br>AAAACGACTTGTCTAAGATGGCAATTGATGGTGATAGAAA<br>CGCGAGGAAAACAATCAATACGCCTCATCGATCAAACTTC<br>TAAAAGCACTTTTTTTGTCGCCGATCAACAACAAATCCTA<br>GAAGTGCTCCCCCGTCAGCAACACTCTGGTACGGCGATC<br>GCCATCGGAAAAAGTCGGGTTGTGCTGTCGTCTCCCCAG<br>CAGGAACTGGATAACTACGTGGGCCTCGGCAAATGCCATT<br>ACCAAGTGCAGCAAAACATTTTCATGCCCCTCAAAGATCC<br>CCAGGCGATCGCCTTTTTAGAAGCGAATCACATTAACCCC<br>AGAGGCCCCCTGGGAAATCGTCGGGGGCTAGTTTTGCGG<br>GAATACATTTGGGTGGCGGGGGATCCGGTTTACATCTATG<br>GCAAGCTGATCCGGCGTGATCGCCAGCACAAAATCCTTAT<br>CCCTTGGGTGATTAGCCACCAACCCCGCGCCCGGATTGTG<br>GTTATTTTGGGGAGTTTGGGGTTAATCGGCCTGAGTTTATT<br>AATTGCAGGGATTACGGTGATTCT |
| NSC11 | FEK30_14770<br>to<br>FEK30_14775 | TTTTCTCCCACAGGATCGTGAATAGATAAACAACGATCAA<br>GAAATAAATTTGTACTGGCATATGATCTTCCCCCACATATC<br>TGAACTAGATGGTGTCTCTCTGTGCTGGACAGAGACAATC<br>AACTGCAGCGAAAAGTTGCATTCCGAAGCACCAACTTTC |

| | | |
|---|---|---|
| | | CCGGTGGTTAATCTTTATGCCACAGGTAAAATTGTTTTTT<br>GTTCCCCATAATCCGAAATTACAACTGGAAATTACGCTAA<br>GTCCTCAGATTTTTCCCAAAGGGATCAACCTGAGACTATT<br>GTTCGCTTTTCTTTACAGCTTTAATATTCCCATTCTTCGCTT<br>TTTACACACACATTCAATGAAAATAAAAGCCAAAATTAGG<br>CCGTATTTTCCCGAACCAATTAAGATCTAAAATAGATCGAC<br>CCACTTAAGGGGGACTCCCCAAGGCAGCACCTAAATGGC<br>ATTCTGCTTGGGCTTAAGGGGTTATTATTCTGTGTAGAAAA<br>CTAAAGTTTCAAAAAATTTAATTATCCCTATCCAATTTCAT<br>CCTAACATCTAACCCCCCCCTCCATTCAAGGGGGAATTTC<br>CAGGGTAAGGTCTTTTGATGGGCTAATGAAACTCATTTAT<br>GCCCACAATGATCACCCTTGTTCCTAGGACAAATATCGTTC<br>GTTTGTACCTAATCATGGGGGTAAATCAATAGATTTTGGCG<br>CGGTTGTCCTGGGCAACAATCCGTAAACTTCTATGGCAGA<br>CCCTGGTTTTTGGAATATTTTCTGTATCATTAAATACAAAC<br>ATTGCAATGATTAGGCTGGCTAATCATCTATGCTCTGTATCT<br>CAAATACTCGTCTCGGAGAAAATCCTG |
| NSC12 | FEK30_15270<br>to<br>FEK30_15275 | ATCCTCCCAGGAAATCCTTAAAACAATCTAAAGAAATTTT<br>TCCTAACCTTCCTTACCCAAGGGAGGTTTTTTATGTGAGTT<br>CACATTTTGTTACGTTACCCAGTCAATACTTGAGCCGCTCA<br>AAAAGTCTGACCTAGAGCAGAAAGTCCCTGAGTATATCG<br>ACTCATTAATCCGGTCTTTCTACTTGGTTTCCTGAGTTGAT<br>TTTCTGCGGAATTTTGGAAATTCAGAGATCTAACCTTAGG<br>GGGAGTCTACTTAAAAACGGCTCTGCTCAACCCTGCAAAT<br>GCCCTACTCTTCTTCTGTCTAGCCCAAGCACTCCCTGAGA<br>AAATTAGCGGCGATCGCCTATAATCATGAAGTTTTGTGAC<br>AGATCTTTTTACAAGATGTAATGTTTAAATGCCGGCAGAC<br>GTTGTATAACATTTACCTAAGATTAAGAGTCACTCGCAGTA<br>CTCCTTAGAAACCCCATAGGTTCCAAGGAACTAGCATGAA<br>CTTTATCTGGCAACTTTGAGAATCTGAGAAATTCAATGAA<br>TGTAAAGTTTCTTAAATGCCAAGGTGAAAAACAAGCAAA<br>AATAGCTGACACTCTTAATTGGCTTTGGGGATTAAGTTTCC<br>AACTCGAAAACAAAACCTTTTATCGACTCTAGGATTTTGT<br>TTTCAGCAAGAGAGCCCCTCAGCACTTGCTTCACTCTTGT<br>TAGTAAGCAAACCGCACAAAATAAATCCCACTCATCAAAA<br>TATAAGTAGGAGATAAAAAC |

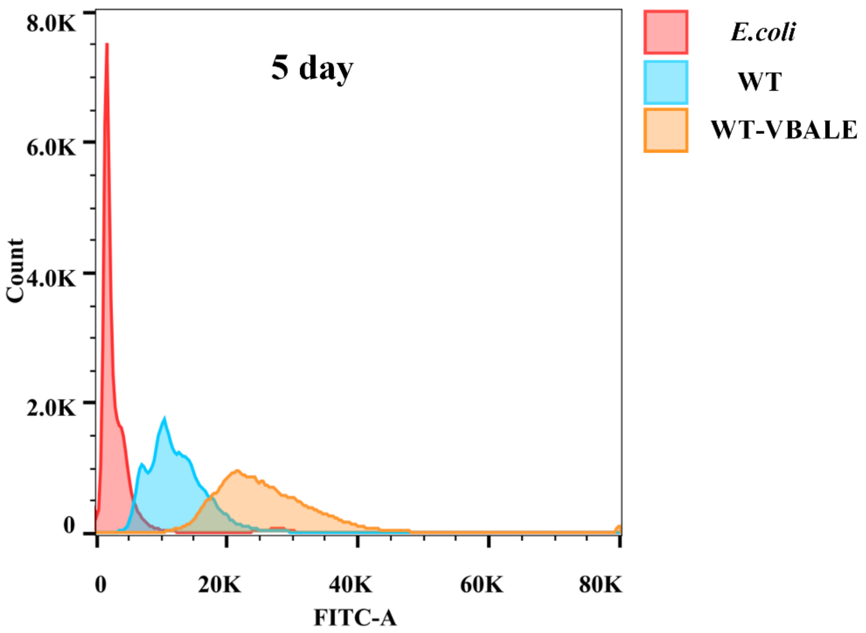


Fig. S1 Genome copy numbers detection of WT and WT-VBALE in the 5th day.

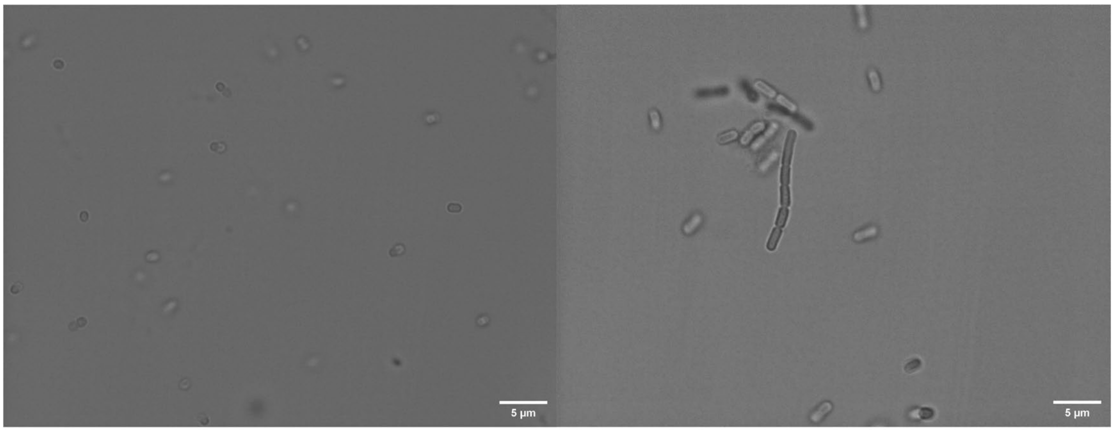


**WT** **WT-VBALE**

Fig. S2 Microscope investigation of WT and WT-VBALE.

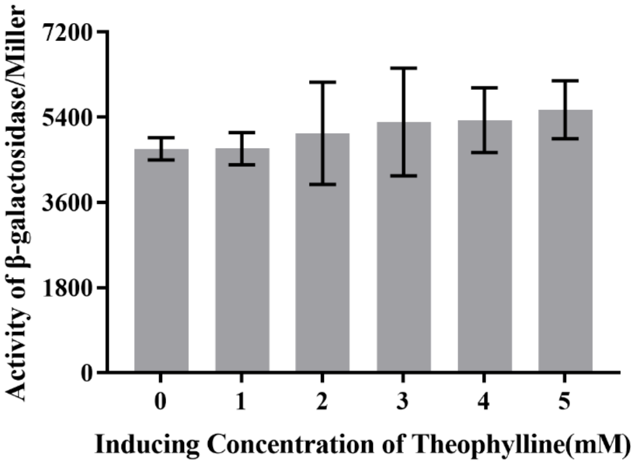


Fig. S3 Induction of LacZ in WT-trc-theoE*-lacZ.
